# AI4Life Open Calls and Public Challenges: why, how, and what we have learned

**DOI:** 10.64898/2026.07.21.739486

**Authors:** Vera Galinova, Mehdi Seifi, Beatriz Serrano Solano, Kristína Lidayová, Damian Dalle Nogare, Agustín Andrés Corbat, Joshua Talks, Edoardo Giacomello, Estibaliz Gómez-de-Mariscal, Mariana G. Ferreira, Caterina Fuster-Barceló, Juan Manuel Battagliotti, Carlos García-López-de-Haro, Benjamin Salmon, Melisande Croft, Si Young Yie, Guillermo Rey-Paniagua, Xiaotian Hu, Sungjun Cho, Aagam Sheth, Chhayansh Porwal, Xiaomeng Li, AI4Life Consortium, Ricardo Henriques, Xinyang Li, Alexander Krull, Anna Klemm, Arrate Muñoz Barrutia, Anna Kreshuk, Wei Ouyang, Florian Jug, Joran Deschamps

## Abstract

Within AI4Life, we ran three Open Calls and three Public Challenges (2023-2025), supporting 22 bioimage analysis projects from 151 applications and engaging 225 challenge participants, with the aim of applying FAIR deep learning in the life sciences. Our experience offers a view of the current state of bioimage analysis, the landscape of available tools, as well as the existing gaps between method developers, tool producers and potential users. It highlights that even after careful selection for AI-ready projects, most still require substantial effort to apply deep learning, and that the field still relies heavily on established, well-rounded methods to solve common problems. We come to the conclusion that for scientific AI in biology, the rate-limiting step is not methods and models but data, annotations, and shared infrastructure underneath them.

## Introduction

AI4Life was a Horizon Europe-funded project (2022-2025) aimed at bringing state-of-the-art AI-based image analysis to life scientists by establishing and supporting services that promote deep learning FAIRness^1,2^. With these overarching goals in mind, the AI4Life consortium conducted various activities, from developing a new format for deep learning models and an accompanying public repository^3^ to organizing workshops promoting tool adoption. Notably, the consortium sought to foster interactions between computational and life scientists, bridging the gap between method developers and those applying these methods. To that aim, we organized three open calls for bioimage analysis projects and three computational challenges.

The Open Calls (OCs) provided life scientists who were facing image analysis challenges with adequate deep learning workflows developed by a team of AI and image analysis experts from our consortium. Selected projects focused on problems that were of interest to the larger community and would therefore impact more than a single research group through re-usability of a solution and open-source tooling. The OCs provided technical support to researchers for a duration of six months, at the end of which the developed pipeline was documented, transferred to the users, and released online.

Technical solutions developed during the OCs were the result of a specific team of analysts’ work and therefore could benefit from further investigation by other experts in the field, either by proposing alternative or superior analysis pipelines. Public competitions have previously proven effective at translating biological questions into well-scoped machine learning problems and driving methodological advances^4–11^. Building on these examples, we organized three Public Challenges (PCs) centered around image restoration, each targeting a different aspect of microscopy image denoising, and running for two months.

In this manuscript, we provide an analysis of the OCs and PCs, describe lessons learned from organizing them, and discuss the pain points encountered along the way. In particular, we believe that our experience reflects the state of the bioimage analysis field and of the interface between method developers and potential users. By summarizing the best practices that we would recommend to avoid the pitfalls we faced, we hope to not only help similar initiatives in the future, but also incentivize the community to continue working towards improved user training, better tool interoperability and FAIRness. A full report and analysis of both the Open Calls and the Public Challenges are provided in the Supplementary Information (SI) for the interested readers. In the following sections, we discuss the main lessons and recommendations that we draw from our experience.

### Open Calls - Lessons learned

The Open Calls were an opportunity to survey the diversity of image analysis problems faced by life scientists, their typical workflows and the current state of the existing solutions offered by the community.

The first challenge was to establish a selection process that would narrow support to projects that were ready to start, solvable within the time frame, and whose solution could be deep-learning-based and transferable to other projects. Our process is shown in Fig. 1, and described in SI section S1.

**Figure 1.**
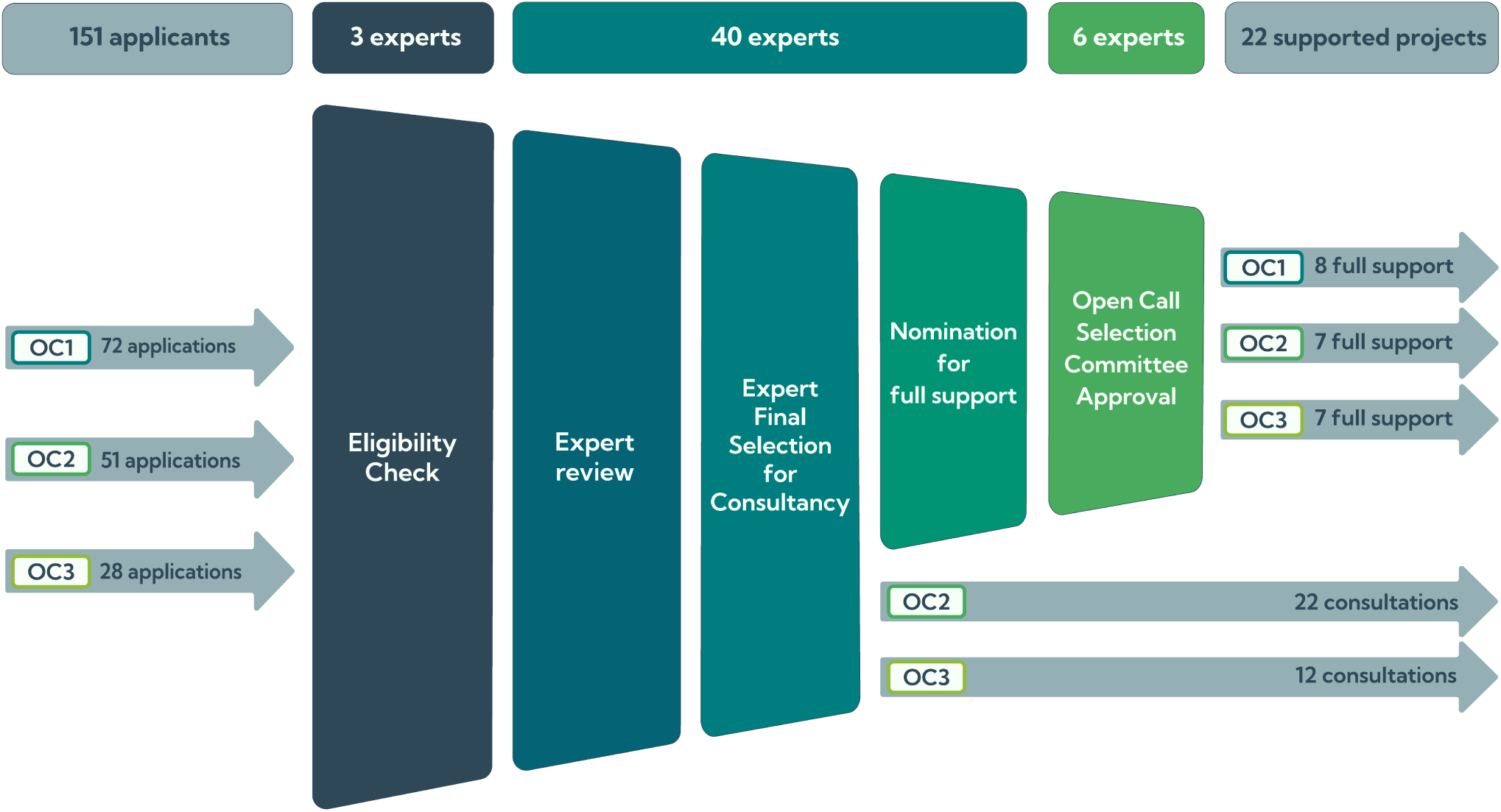
Selection procedure and analysis support workflow for the three Open Calls. The number of experts involved in eligibility checks, reviewing applications, consultation calls, and members of the selection committee is indicated in the top row. OC 1 did not include a consultation step.

Across the three OCs, we progressively moved from pure remote scoring to interactive expert dialogue as OC 1 revealed a high variance in both data readiness and the actual relevance of deep learning to the proposed problem. Multiple projects were ultimately solved with classical image analysis or existing tools, including pre-trained deep learning models, while others were delayed by data transfer or the need to re-acquire data.

To reduce this mismatch, OC 2 and 3 introduced three decisive changes: mandatory upload of representative data, a template-based application, and a consultation round with experts. The consultations required substantial additional work, but were the most reliable way to identify ready-to-start projects, re-frame over-ambitious ones, and give useful feedback even to non-selected applicants. Finally, the reviewing scores were simplified to only 3 categories to increase the transparency of the scoring: readiness, applicability of deep learning, and scientific excitement (SI section S1.2).

Challenges faced during the OCs are familiar to image analysis facilities and highlight the need to build better community resources to educate users about the availability of user-friendly tools. As described in SI section S2.3, we observed that many widely used tools still failed to get into the hands of users (e.g. Cellpose^12^, CellProfiler^13^, QuPath^14^, ilastik^15^, Labkit^16^, or the napari^17^ ecosystem), while users rely heavily on proprietary software, hindering the spread of reproducible analysis and FAIRness. The nature of microscopy images and scientific goals often leads to complex pipelines, chaining several existing tools and requiring custom scripting, as exemplified by the pipelines developed for the OC (see SI section S2.5).

The recurrence of specific tools in the developed pipelines demonstrates that these are cornerstones of bioimage analysis. Notably, this was the case of Cellpose^12,18^ and StarDist^19^ for instance segmentation, and Noise2Void for denoising^20^. Similarly, while the field is moving heavily towards Python under the impulse of scientific Python libraries and deep learning, ImageJ2/Fiji^21,22^ remains an important ecosystem. Several ImageJ2/Fiji plugins provide solutions to widely encountered challenges and are accessible to non-specialists (BigWarp^23^, Labkit^16^, TrackMate^24^). Well-established tools often owe their success to both effectiveness and user-friendliness^25^. State-of-the-art methods, on the other hand, may suffer from complex installations and use. Newer architectures (transformers, foundation models, diffusion-based methods) are absorbed by the

field selectively, and the OC pipelines suggest the absorption criterion is whether the new method reduces the data and annotation burden on users rather than whether it improves a benchmark by a few points. This is exemplified by the success of SAM/SAM2^26,27^ in segmentation, which has led to a sharp increase in segmentation tools relying on it as a pre-trained model. Such tools have been used during the OCs, or developed for an OC project^28^, hinting at a new direction for tool development in the era of large foundation models developed outside of the field of bioimage analysis.

Another issue apparent from both applications and consultations is the need to spread realistic expectations about the capabilities and requirements of deep learning algorithms^29,30^. Two different aspects of deep learning in particular appear to be generally misunderstood: the quality of annotations required to train a deep learning model, and the current capacities of deep learning methods. For the latter, the consultation round was instrumental in managing user expectations and re-framing the scope of the projects based on more realistic goals. Data labeling, on the other hand, is a time-consuming activity and labels quality is often overestimated by users.

Only 8 out of 22 selected projects were fully ready at the start of the support period, with complete, usable data and sufficient annotations available to start working on the project. Even more striking, only 2 of them had sufficient ground-truth data to directly envision using deep-learning as a first step (see SI section S2.6). In projects that required substantial data labeling, the analysis pipelines often made use of alternative methods or attempted to leverage sparse labeling (see SI section S2.5).

While some projects turned out to be more challenging than expected, most of the incurred delays were a result of difficulty in exchanging data, in particular large datasets. Often, data exchanged relied on private services, external to the scientific institutions (see SI section S2.6). Not only does this present a security risk, but it also highlights the lack of shared data infrastructure across European public research institutions and the dissemination of local solutions.

Furthermore, while most applicants agreed in principle to publicly release their data, the immediate benefit of doing so is rarely visible to the user when sharing has to happen, typically near the end of a project, when attention has already shifted elsewhere. The consequence is that data release is consistently deprioritized: of 22 supported projects, only 17 had data deposited in a public repository by the time of writing, with 1 still pending and 4 unable to contribute data at all (see SI section S2.6). Furthermore, most of the projects incurred significant delays in releasing data publicly, with 11 projects having data uploaded 6 months or more after the official end of the project and an overall average of about 8 months delay between end of support and public data release. FAIR data sharing is not a one-time act at publication but a sustained effort that extends well beyond the visible end of a project, and current incentive structures do not reward that effort. Concretely, we found it more effective to identify shareable subsets early, begin the upload process before the analysis pipeline was finalized, and treat the data deposit as a parallel deliverable rather than a closing step. Even with this strategy, attrition was substantial.

The deeper issue underlying low data-sharing rates is not unwillingness but invisibility of return. Researchers who share image data rarely see direct credit for it: citations accrue to method papers and biological findings, not to the underlying datasets that made either possible. As long as this asymmetry holds, FAIR data sharing will remain something individuals do despite their incentives rather than because of them. Echoing others^31^, we believe that in order to meaningfully shift this trend, two main avenues should be pursued. First, training: most life scientists encounter FAIR principles as compliance language rather than as a working description of how reusable data is structured, annotated, and deposited^32^, and the gap between willingness to share and data usable by others is wider than most applicants realize at submission time. Second, recognition: dataset publications, repository deposits, and reusable annotations should count toward hiring, promotion, and funding decisions on a comparable footing with method publications. Neither change is within the power of any single project, but both are within the control of the community that funds, hires, and evaluates its members.

Taken together, the lessons from three Open Calls describe a community that is technically capable, intellectually engaged, and culturally aligned with open science, but is operating under a certain lack of infrastructure, training resources, and shared norms that AI methods quietly assume.

### Public Challenges - Lessons learned

The Public Challenges offered a complementary perspective, producing fixed benchmarks for method developers to explore diverse solutions.

The central lesson from the Public Challenges is that the bottleneck is not community interest in biological imaging, but the rarity of problems that are immediately benchmark-ready. Our initial vision was to draw challenge tasks directly from the Open Calls, but in practice this proved considerably more difficult than anticipated. Many datasets lacked the quality or quantity that deep learning benchmarking requires. Beyond data readiness, many proposals were either too narrowly scoped within a specific biological subfield to attract broad computational interest, or described multi-step experimental workflows that were difficult to abstract into a single, well-defined task, which is a prerequisite for any accessible and fairly evaluated challenge.

This gap motivated us to shift toward publicly available datasets and synthetically generated data as the foundation for challenge design (see SI section S3). While no dataset perfectly captures the diversity of real-world biological imaging, curated public resources offer a practical and transparent starting point: data provenance is known, baselines can be established, and participants can access training data openly. This approach also naturally promotes reproducibility. Future challenge organizers in the biological imaging space should treat data readiness as a central criterion, and the availability of suitable public datasets as a precondition rather than an afterthought.

The three challenges focused on image denoising in three different settings (see Fig. 2): unsupervised (PC 1), supervised (PC 2), and domain-specific calcium imaging (PC 3). Together, they provided a useful snapshot of which methods participants currently reach for under different training regimes (see SI section S4). In all three settings, participants largely relied on established pipelines: Noise2Void and its variants were the most frequent submissions, reflecting the method’s wide adoption and ease of deployment. In the unsupervised setting (PC 1), however, COSDD^33^ placed first on three of the four leaderboards and a narrow second on the fourth, where an N2V submission edged it by 0.01 dB. This highlights the potential of methods that have not yet reached mainstream adoption. The supervised challenge (PC 2) attracted more methodological diversity, with participants adapting U-Net architectures and also trying generative adversarial networks (NAFNet-GAN^34^), the latter topping three out of four leaderboards, suggesting that supervised denoising is both more familiar and more open to experimentation. PC 3, focused on the more demanding calcium imaging setting, showed the slowest submission rate. Here, a transformer-based model (Restormer3D^35^) topped all four leaderboards, while a well-tuned N2V 3D baseline also remained highly competitive and consistently placed in the top three. Taken together, these results suggest that established methods remain the default choice for participants, but that alternative architectures can become clearly advantageous in settings that are technically more demanding.

**Figure 2.**
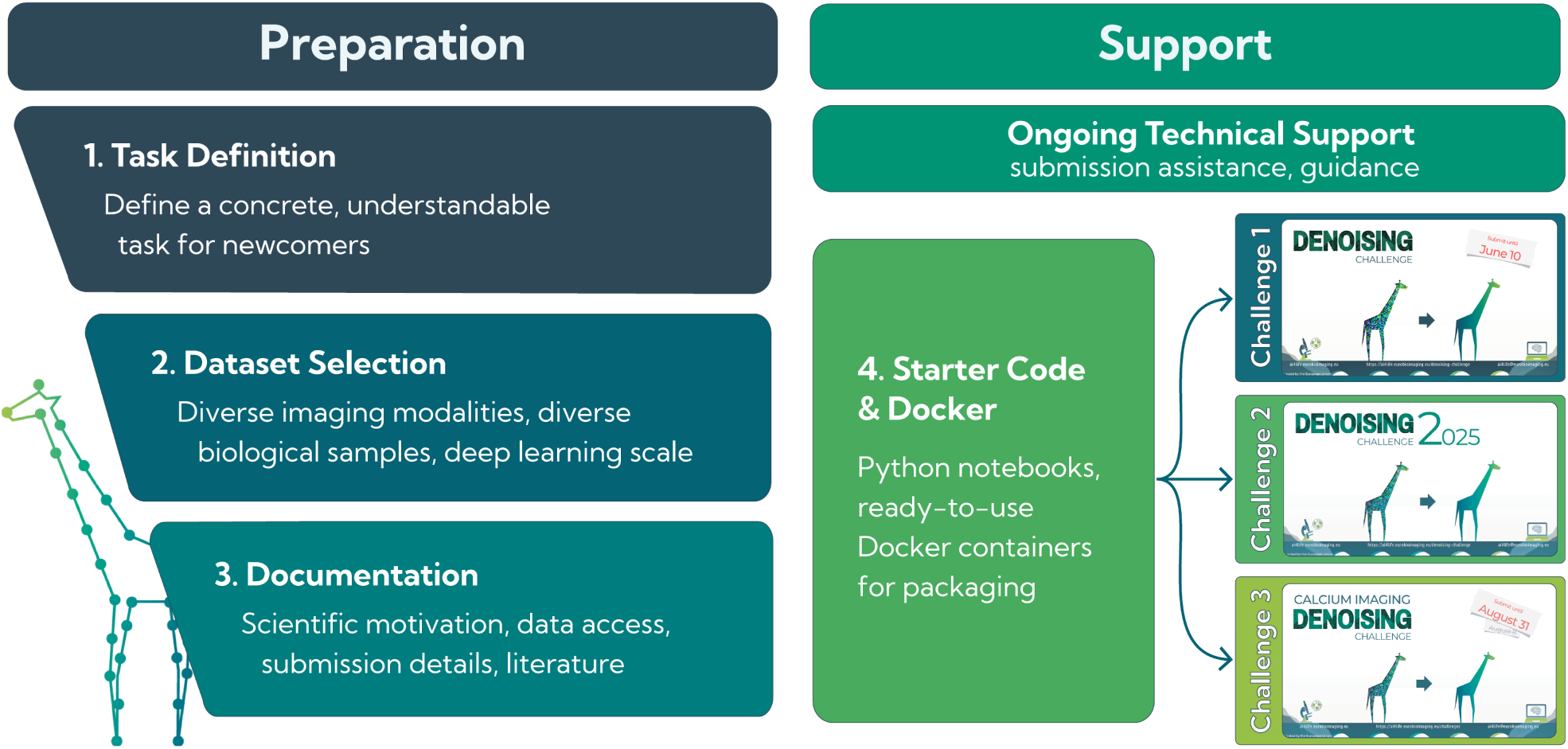
**Challenge organization process.**

Running these challenges also made clear that community benchmarking reveals performance differences that are hard to see from individual method papers alone: a method may be well-known, yet its relative strength across diverse datasets and noise regimes only becomes apparent through direct comparison^30^.

The geographic diversity of registrants was an encouraging sign of broad community interest (see SI section S4). However, the overall number of successful submissions remained modest compared to established challenges in computer vision or medical imaging. We attribute this primarily to the absence of monetary or conference-based incentives: many prominent challenges offer cash prizes, hardware, or co-location with major venues, sometimes accompanied by paper authorship for top participants. Without such incentives, participation is driven almost entirely by intrinsic scientific interest, reaching a narrower audience. This suggests that future challenges in biological imaging would benefit from strong incentives, such as prize money, hardware, paper authorship for top participants who release their code, broader outreach through community channels such as the image.sc forum and social media, and closer alignment with conferences or journals to make participation more accessible and attractive to a wider community.

The three Public Challenges revealed that we are constrained by the scarcity of benchmark-ready biological problems, openly reusable datasets, and the incentives needed to sustain broad participation.

## Discussion

What emerged from both initiatives was not two separate sets of issues, but the same underlying bottleneck viewed from two different angles. Three years of structured support for 22 bioimage-analysis projects, alongside three public denoising challenges, gave us an empirical view of the gap between the methods the field has published and the conditions under which those methods can actually be applied in practice. That gap is the central finding of this work. The algorithms exist, in many cases as community-supported, sometimes as well-engineered tools, but data are too small, too coarsely annotated, too poorly standardized, too costly to share, and too fragmented across institutional infrastructures.

The Open Calls showed that this gap can be narrowed project-by-project when expert support is available. 18 of 22 projects produced reproducible, openly released analysis pipelines, suggesting that targeted bioimage-analysis support can generate durable value even when starting conditions are uneven. The Public Challenges, by contrast, showed how difficult it remains to convert real biological needs into benchmarks that are sufficiently well scoped, well resourced, and broadly accessible to invite community-wide comparison. Together, these two observations sharpen the same point: methodological progress alone does not translate into practical impact unless the surrounding data and infrastructure are mature enough to carry it.

These observations are easier to dismiss in isolation than together. We observed the dominance of TIFF and proprietary microscopy formats over next-generation file formats (8 of 151 applications), the persistence of proprietary analysis software despite mature open alternatives, the fragmentation of data transfer across 14 different services (for 22 projects), and the fact that many well-adopted open tools remain off the radar of the researchers who would most benefit from them. Each of these is, on its own, a manageable inconvenience. Together, they describe a field whose data infrastructure, tool ecosystem, and cultural norms remain some distance from the level of maturity that current AI discourse often assumes.

We do not see this as cause for pessimism. Frontier AI systems trained on web-scale text were possible because the training data was already there. Our field has none of those conditions fully developed and is now building them deliberately with the examples of the Image Data Resource^36^, the BioImage Archive^37^, the BioImage Model Zoo^3^, OME-NGFF^38^, community standards^2,32^, and projects such as AI4Life itself. What this work shows is how far that building still has to go, and how concretely it can be done. The community resources required are not exotic: shared data-transfer infrastructure with institutional support, sustained annotation efforts on biologically meaningful datasets, training material that calibrates user expectations of what deep learning can and cannot do, and incentive structures that reward dataset publication on par with method publication. None of this is a research problem in the conventional sense. It is infrastructure work, dispersed and undramatic. It is also the rate-limiting step on what scientific AI in biology will deliver in the coming decade^30^.

### Recommendations

If these constraints are largely infrastructural, the response must also be practical. The lessons we draw from three Open Calls and three Public Challenges therefore fall into actionable recommendations, grouped here by the audience best placed to act on them.

#### For life scientists submitting projects

Deep learning generally benefits from very high quality labeling. Before entering any AI collaboration, it is helpful to keep in mind that labels likely need substantial revision or extension. Plan annotation effort early in the discussion, and treat data and annotations as one of the slowest-moving parts of any AI collaboration.

When possible, adopt next-generation file formats^38^ for projects involving volumetric, multichannel, or large-scale imaging. TIFF and proprietary formats remain workable for small data, but can make downstream reuse and scaling more difficult.

Explore the open-source bioimage-analysis ecosystem (ImageJ2/Fiji^21,22^, CellProfiler^13^, QuPath^14^, Cellpose^12^, ilastik^15^, Labkit^16^, napari^17^, etc.) before commissioning custom development. In many cases, a method that addresses much of the problem already exists and may be within reach with some guidance or training. The bioimaging community will be happy to help you find a good solution on the Image.sc forum^39^, and the answers will also help others.

Finally, it helps to approach deep learning with realistic expectations: it is not a generic intelligence applied to images, but a class of methods with concrete data and annotation requirements^29,30^. In our experience, projects are much more likely to progress smoothly when those requirements are made explicit from the outset.

#### For image analysts carrying out projects

Verify data and annotation readiness before scoping methods. The Open Call experience showed that remote scoring alone is often not enough to distinguish ready projects from those that still need some preparation. A structured consultation with sample data in hand was the most cost-effective filter we found.

Expect to spend a meaningful fraction of any project on data transfer, format conversion, and annotation triage. Budget for it explicitly.

Default to FAIR pipelines^32^ from the first commit, not as a publication afterthought. Data uploads after the fact are the most common point of attrition. Where possible, prefer the tools that are the best fit for the project, including when they are not the ones you developed yourself. Conversely, recognize that some tools and methods (Cellpose^12^, StarDist^19^, Noise2Void^20^, ilastik^15^, Labkit^16^) are cornerstones for good reason and are often worth considering before custom development.

Finally, team up with others. There are many opportunities, and meetings such as “I2K: From Images to Knowledge” or image analysis networks such as GloBIAS^40^ are a good way to strengthen the international analysts community.

#### For bioimage-analysis facilities and community organizers

Many widely-used tools still do not reliably reach the users who would benefit from them most, suggesting that the main gap is one of discoverability and training rather than methods alone.

Maintain community-driven, regularly updated documentation of what current tools can do, what data they require, and where they fit into typical pipelines.

For open calls or similar support programmes, include sample data in applications, use a fixed presentation template for project descriptions, and run at least one consultation round before final selection.

Keep scoring schemes simple and interpretable, ideally with three or four categories that can be discussed transparently. Identify two or three preferred data-transfer channels by data size (with institutional support where possible, e.g. Globus^41,42^), and start the data-release process before the pipeline is finalized rather than after.

In our experience, these changes substantially improved selection accuracy and reduced project attrition between OC1 and OC3.

#### For method developers, research software engineers and challenge organizers

State-of-the-art methods are most useful to biology when they are also installable, runnable on real data, and accompanied by clear guidance on when they apply.

The success of Cellpose^12^ and Noise2Void^20^ owes as much to their accessibility as to their algorithmic merits. Conversely, the rapid uptake of SAM2^27^ derivatives in our pipelines shows that the field will absorb foundation models quickly when they reduce the data burden for users.

When releasing a method, consider what other researchers would need in order to apply it to their own data, and take the extra steps needed to make that process as easy as possible^29,43^. If developing a tool, ensure accessibility, documentation and interoperability with existing ecosystems^25,44^.

For benchmarking, treat data readiness as a precondition rather than an afterthought: many real biological problems do not yet naturally fit the shape of a well-posed challenge, and curated public datasets therefore remain the most viable foundation. Future challenges in this space will benefit from stronger participation incentives. This includes prize money, hardware, paper authorship for top entries, and closer alignment with conferences and journals to reach beyond the existing community.

#### For funding bodies and institutions

The most consistent finding across both Open Calls and Public Challenges is that the limiting resource is shared infrastructure. Recognize fragmented data transfer as an infrastructure gap rather than an individual failure.

Invest in cross-institutional data-transfer infrastructure, in public bioimage repositories^36,37^, in annotation efforts on community-prioritized datasets, and in the open-source tool maintainers who underpin the entire ecosystem.

Treat this work as strategic capacity building for scientific AI in biology, not as a peripheral service layer. The community is already building these resources^45^, but the question is whether funding structures will recognize them as the frontier work that they are.

## Acknowledgements

We would like to sincerely thank the bioimage analysis community for its continuous development of open tools and support via the Image.sc forum. We are grateful to all applicants and participants involved in the Open Calls and Public Challenges initiatives, and to the reviewers involved in the Open Calls: Aitor Gonzalez-Marfil, Anil Kumar Mysore Badarinarayana, Anirban Ray, Ashesh, Ben Woodhams, Callum Tromans Coia, Daniel Sage, Dominik Kutra, Elena Buglakova, Gisele Miranda, Kate Gill, Mario Del Rosario, Martim Gamboa, Martin Schätz, Marvin Albert, Pablo Rodríguez Pérez, Thibault Lagache, Varun Kapoor, and Weize Xu, and the participants to the Solvathon Hackathon organized at EMBL: Fynn Beuttenmüller, Jonas Hellgoth, Qin Yu, Craig Russell, Constantin Pape, Anwai Archit, Fariha Mahzabin Annesha, Ashesh, Igor Zubarev, Ankit Roy, and Sushmita Nair. We also thank the Bioimage Analysis Unit, National Facility of Data Analysis and Handling, Human Technopole, and the BioImage Informatics Unit, SciLifeLab, for support, feedback and constructive discussions. We also thank Matthew Hartley, Aybuke Kupcu Yoldas, Teresa Zulueta-Coarasa and the whole BioImage Archive team for their support in uploading datasets. We are grateful to all participants in the Public Challenges, and in particular to Kino Sun (user a897574323) and Yuanchen Wang (user wangyuanchen) for placing in the top 3 and making their training code publicly available.

## Author contributions

The Open Calls were organized by VG, MS, BSS, DDN, FJ and JD. OC projects were supported by VG, MS, KL, DDN, AAC, JT, EGdM, MGF, CFB, JMB, CGLdH, and JD, with supervision from EGdM, RH, AK, AK, AMB, AnK, WO, FJ, and JD. Challenges were organized by VG, EG, XinL, AlK and FJ, with support from BSS, AMB, and WO. BS, MC, SYY, GRP, XH, SC, AS, and CP participated in the public challenges, achieved top-3 placements on the leaderboards, and contributed their training code for reproducibility, with BS’, MC’s, GRP’s, and SC’s participation supervised by AlK, JD, AMB, and XiaL, respectively. Analysis of OCs and PCs meta-statistics were carried out by VG, MS, BSS and JD. The manuscript was written by VG, MS, BSS, FJ and JD with inputs from co-authors.

## AI4Life Horizon Europe Programme Consortium

## Supplementary Information

### S1 Open Calls organization

#### S1.1 Process description

In this section, we describe the process organization towards which we converged after three OCs. The OCs ran for three consecutive years (2023, 2024, and 2025), following similar structures: (1) application step, (2) reviewer recruitment, (3) reviewing process, (4) project ranking, (5) consultation and project selection, and (6) project support. This workflow aimed at identifying, evaluating, and supporting bioimage analysis projects suitable for AI-based solutions. The process was designed to balance accessibility for life scientists with rigorous technical assessment, while ensuring efficient use of expert resources. It combined structured applications, anonymised review, expert consultation, and hands-on project support, with an emphasis on project readiness, feasibility, and reusability of outcomes. The structure was continuously refined and adjusted throughout the calls based on the previous year experience, leading to a process organization that can serve as a reference for future open calls. For each selected project, a team of deep learning and image analysis experts built a suitable pipeline to solve the challenges faced by the researchers, pipeline which was then released online according to the FAIR principles together with the data used in the project.

##### Application

In the first step of the process, applicants submitted a standardized application consisting of a brief project description and representative image data, including annotations when available. See Appendix A1 for OC3 application form. A common presentation template of four slides and an example project were used to ensure consistency across applications and to facilitate rapid assessment of project goals, data characteristics, and existing workflows by the reviewers (see Fig. S1). Guidance was provided on terminology and ground truth quality with an illustrative figure in the application process, helping applicants describe their data and challenges accurately, and improving the reliability of reviewer assessments. All applications underwent an initial pre-screening by the main organizers to exclude submissions that were incomplete or insufficiently documented. This step focused on the clarity and completeness of the application, ignoring scientific merit, and ensuring that only projects suitable for external review progressed further.

**Figure S1.**
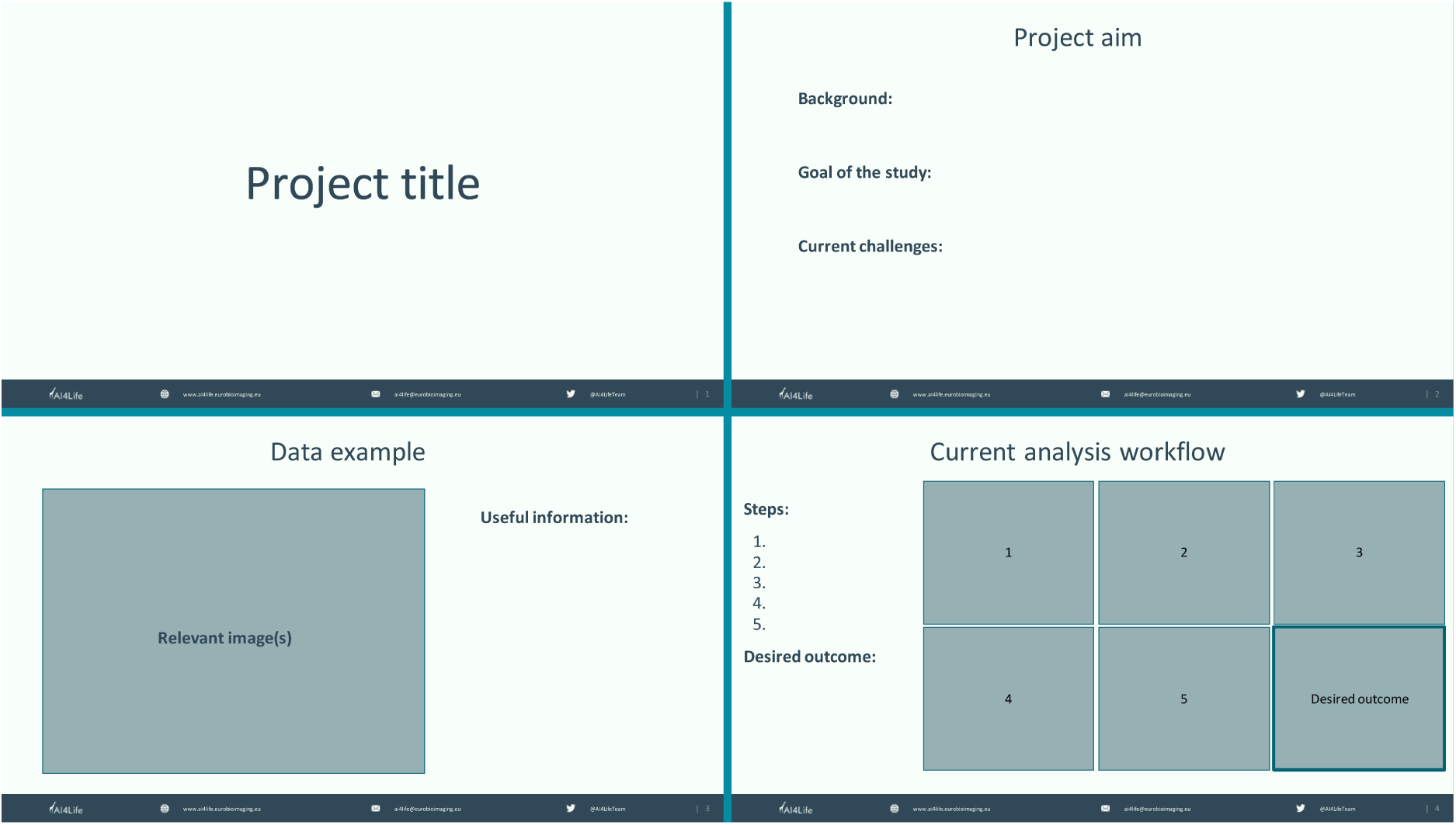
Project template for applications in OC 2 and 3. Each application was requested to upload a presentation in the format of their choice (.pdf, .pptx, .key, etc.) following this template to standardize the information reviewed by reviewers.

##### Review

During the application window, we recruited reviewers both internally to AI4Life and in the community of bioimage analysts. Advertisement for registration as reviewers were sent by emails to image analysis facilities, posted at conferences (e.g. I2K) and advertised on the image.sc forum^1^. Once the pool of reviewers was assembled, eligible project applications were evaluated through anonymized peer review using predefined scoring criteria. Reviewers were guided to assess project readiness, applicability of deep learning, feasibility within available effort, and scientific excitement, and suitability for the aim of the OCs (see Section S1.2). Measures were taken to prevent reviewing errors, including controlled project assignment and consistency checks, and to identify systematic reviewer bias. Review scores were aggregated using a simplified and automated scoring scheme to generate a ranked list of applications. A score cut-off was applied to shortlist projects moving to the next selection phase.

##### Consultation phase

In OC 2 and 3, shortlisted projects were invited to a consultation phase consisting of discussions with a panel of image analysts and AI experts. These consultations were used to verify data readiness, clarify technical challenges, identify appropriate tools, and assess feasibility. When necessary, a second consultation was organized to allow the panel of experts to come to a shared understanding of the project’s current state. The consultation phase panels were composed of members of AI4Life partners’ institutes and external image analysts. We ensured that typically three or four experts were present for each session, with the aim of pairing expert domain knowledge with relevant projects.

##### Project selection

After the consultation phase, all experts who had participated in the various meetings with the applicants assembled and discussed the projects. Written feedback from the experts informed the final selection, which was performed by adjusting the technical feasibility and readiness scores from reviewers. Top-scoring projects were selected in order up to the maximum capacity for support, and in compliance with the requirements of transnational access imposed by the Horizon Europe funding.

##### Project support

Selected projects entered a support phase formalized through signed agreements. Data transfer strategies were defined early to avoid delays, and open data release plans were developed in alignment with FAIR principles. AI4Life experts provided hands-on support to develop reproducible, open-source analysis pipelines. Progress was monitored through regular meetings that addressed data acquisition, labeling needs, pipeline usability, and coordination between experts, with early user involvement to facilitate adoption beyond the support period.

##### Project conclusion

At the end of each project, a description of the analysis pipeline was uploaded to GitHub and Zenodo, alongside any custom code or tool developed for the need of the project. The GitHub repositories served as basis for transferring the pipeline to users. When possible, data used in the project was also released on the BioImage Archive^2^. Finally, all compatible fine-tuned or de-novo trained deep learning model were saved in the BioImage Model Zoo format and uploaded to the BioImage Model Zoo repository^3^.

#### S1.2 OC3 project ranking

We appointed an international team of expert reviewers who helped us to evaluate project applications according to the following criteria: readiness of the project, applicability of deep-learning and excitement about the project. Reviews were sent via a form (see Appendix A2).

All OCs had different criteria, here we only describe OC3 ranking process as these were the criteria to which we converged.

The following criteria were aggregated across all reviewers for each project.

1. Average readiness: *µ_r_ ∈* [0, 5]
2. Average DL applicability: *µ_dl_ ∈* [0, 5]
3. Average Excitement: *µ_ex_ ∈* [0, 5]
4. Sum of AI4Life relevance: *s_ai_ ∈ {*0, 1, 2*}*
5. Sum of Consultation relevance: *s_co_ ∈ {*0, 1, 2*}*

A general score was computed as the sum of the reviewing criteria weighted by the reviewer’s belief that the project was a good AI4Life project or would greatly benefit from consultation:

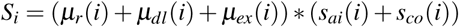

The projects were ranked by their *S* score and projects qualifying for trans-national access (as per the Horizon Europe rules) were selected for consultation. Projects not qualifying for trans-national access were selected up to the quote allowed by the funding constraints.

#### S1.3 Methods

The entire Open Calls process was carried out using Google Forms for the project applications, applications to be a reviewer, and review submission. Applications for supports were saved with a unique ID in a Google Drive folder accessible only to the main organizers. The data was automatically anonymized with an Apps Script and copied to a reviewer-only accessible folder. Each reviewer had only access to the relevant project applications. Scores were aggregated in a Google Sheet, in which the processing was performed.

### S2 Open Calls outcomes

#### S2.1 Applications statistics

The OCs received 151 applications throughout the three calls (OC 1: 72, OC 2: 51, and OC 3: 28) from 28 countries worldwide, spanning all inhabited continents. Most submissions originated from European institutions, particularly from Germany (27 applications), Portugal (20), Italy (14), France (10), and the United Kingdom (10) (see Fig. S2a), while substantial participation was observed from the United States (16). Of these 151 applications, 32 projects underwent consultations (OC2: 22, OC3: 12) and 22 were ultimately selected for support (OC 1: 8, OC 2: 7, and OC 3: 7). Due to transnational access constraints from the funding body, the number of consultations and project supports was capped for non-EU countries. The distribution of home base countries for the selected projects was coherent with the overall applications (see Fig. S2a, light bars). EU countries with the most selected projects were Germany (6), France (3), Portugal (2), Italy (2) and the Netherlands (2). Three projects from non-EU countries were selected, originating from the United States (2), Australia (1) and Canada (1).

**Figure S2.**
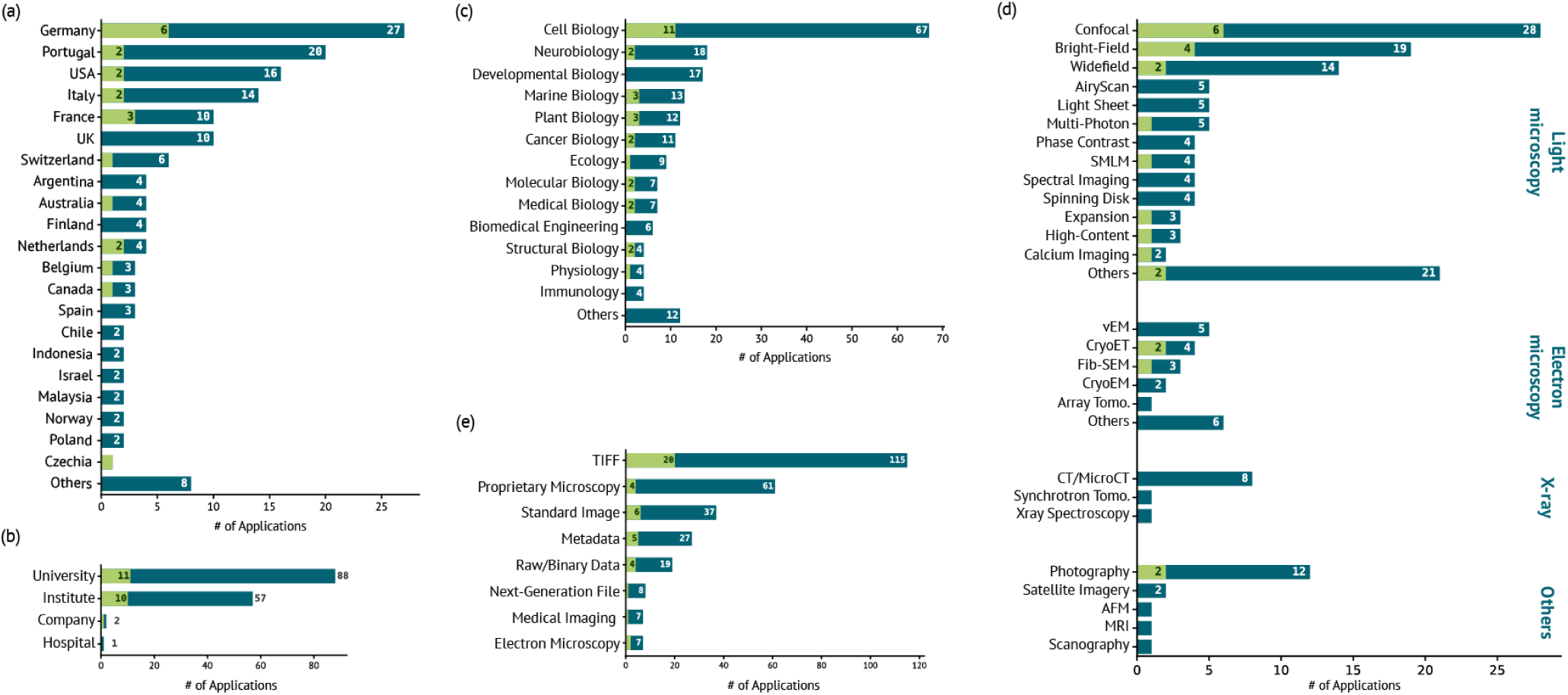
(a) Countries in which applications (dark) and selected projects (light) home institutions are based. Others correspond to single applications from Austria, Denmark, Hungary, Mexico, Serbia, Slovakia, South Africa, and Sweden. (b) Type of organization from which applications (dark) and selected projects (light) originated. (c) Scientific domains of applications (dark) and selected projects (light). Note that each application could declare multiple scientific domains. Others correspond to Agronomy, Biophysics, Genetics, Histology, Morphology, Pathology, Remote Sensing, and Virology. (d) Imaging methods of the applications (dark) and selected projects (light) by categories: light microscopy (LM), electron microscopy (EM), X-ray, and others. “Others” category in EM and LM includes projects for which the particular method was not explicit (EM and LM), or was mentioned by a single application only (LM). “LM Others” includes STED, OCT, ODT, and flow cytometry (selected project). Selected CLEM project was counted in both LM (wide-field) and EM (Others). (e) Data formats, as declared during applications (dark) and for the selected projects (light).

Applicants showed a balanced gender distribution (statistics only collected during OC2 and OC3), with a slight predominance of female applicants (47 females, 30 males), while a small number reported non-binary gender or preferred not to disclose (2). Applicants represented a wide range of nationalities (see Fig. S3), predominantly European (53 out of 79 applicants). Note that nationality and gender information were not collected for Open Call 1.

Applicants home institutions were overwhelmingly universities (88 applications, of which 11 were selected) and research institutes (57 applications, 10 selected), as shown in Fig. S2b, and few applications came from companies (2 applications, 1 selected) or hospitals (1 application).

Submitted projects covered a broad range of scientific domains, with a strong concentration in cell biology and adjacent fields. The most frequent domains declared during applications (see Fig. S2c, dark bars) included cell biology (67 applications), neurobiology (18), developmental biology (17), marine biology (13), plant biology (12), and cancer biology (11), reflecting both core bioimaging communities and applied life-science use cases. Less frequent submissions spanned specialized areas such as structural biology, physiology, or immunology. Note that applications could declare multiple scientific domains. Once again, selected projects were sampled appropriately from these domains (see Fig. S2c, light bars), with a majority of them belonging to cell biology (11 projects), and spanning most declared scientific domains. Structural biology, while being declared by only 4 applications, was well represented with 2 selected projects.

**Figure S3.**
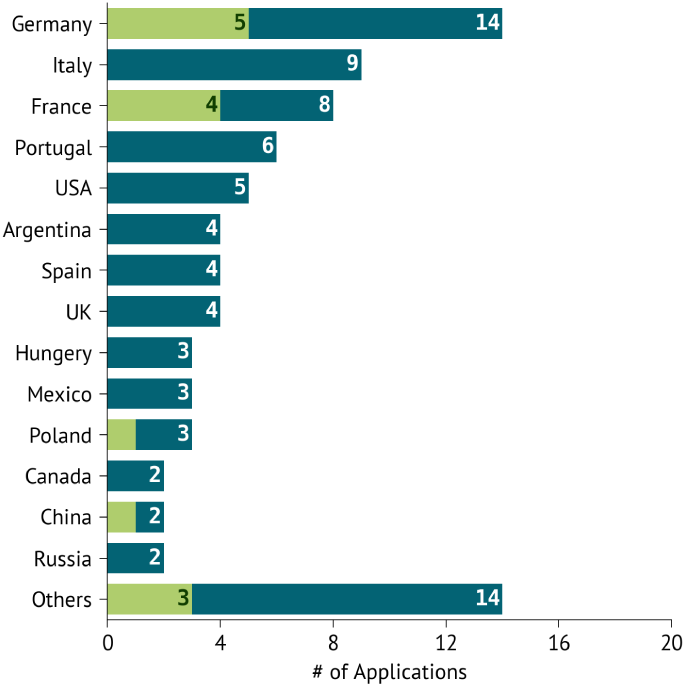
Self-declared applicant nationalities collected during OC 2 and 3. “Others” category consists of Chile, Croatia, Czech, Greece, Indonesia, Ireland, Israel, Japan, Malaysia, Netherlands, New Zealand, Sweden, Switzerland, and Turkey. EU nationals accounted for 53 out of 79 applicants (some applicants were multinationals).

#### S2.2 Data statistics

We extracted the source imaging approaches from the applications, with applications potentially declaring multiple ones. Applications were predominantly using light microscopy (121 applications, see Fig. S2d), far more than electron microscopy (21) and X-ray (10). Other types of techniques (17 applications) included natural images and photography (12), mostly by projects from ecology, plant biology, and agronomy domains, and few applications with satellite images (2), Scanography (1), AFM (1), or MRI (1). Within light imaging, widespread imaging methods dominated the applications, with confocal (28 out of 121 applications), bright-field (19), or widefield (14). More recent and specialized techniques, such as Airyscan, light sheet, multi-photon, or single-molecule localization, were used by a few applications (4 to 5 applications). The rest of the methods covered a wide range of the microscopist toolbox. For many applications, the particular imaging method was not explicit and was therefore grouped in the “Others” category in light microscopy, together with single occurrences (STED, OCT, ODT, flow cytometry). Electron microscopy methods covered the most common methods available to scientists (e.g. vEM, cryoET, cryoEM, FIB-SEM). Several applications only mentioned scanning or transmission electron microscopy, and were grouped together with undefined EM methods (“Others”, see Fig. S2d). X-ray imaging or spectroscopy was largely dominated by computed tomography, with 8 applications out of 10. None of the X-ray applications was selected.

For all applications, applicants could declare image dimensionalities and data formats. There were slightly more applications declaring 2D (99 out of 151 applications) than 3D (83). Multi-channel imaging concerned 63 applications, and time-series were limited to only 40 applications out of the total. Note that, here again, each application could select multiple categories.

Finally, TIFF format was by far the most common declared data type involved in the application (115 out of 151 applications, see Fig. S2e), before proprietary microscopy formats (61), or standard non-scientific image formats (37). Next-generation file formats, such as HDF5 or Zarr, were declared in only 8 applications in total. Here again, multiple options could be selected by each application.

#### S2.3 Tool statistics

Applicants had the opportunity to mention the tools that they tried or use for image analysis in the context of their project. While in OC 1 tools were mentioned as free text, we added tools and tools categories suggestions to OC 2 and 3 (see section A1). In Fig. S4a, we aggregated the answers from all OCs into the categories used in OC 2 and 3. As expected from a call dominated by life sciences applicants, ImageJ2/Fiji^4,5^ was the most cited tool (116 out of 151 applications). Scripts (“ImageJ macro, Python, R, Matlab, etc.”) were cited in 60 applications. Note that ImageJ2/Fiji macros were categorized in scripts, thus underestimating the use of ImageJ2/Fiji by the applicants. The next category was commercial analysis software (“Zeiss, Arivis, Imaris, Amira, Huygens, etc.”) with 41 applicants declaring using them. Additional commercial tools used for statistical analysis and plotting, namely GraphPad Prism and Excel, were mentioned in 25 applications. Specialized libraries for machine learning (PyTorch, TensorFlow, scikit-learn, etc.) were cited 19 times. Interestingly, popular tools in the image analysis community, such as ilastik^6^ (15), Cellpose^7,8^ (14), napari^9^ (14), CellProfiler^10^ (12), QPath^11^ (8), Labkit^12^ (4), or Icy^13^ (2), were cited less than 15 times each. Tools allowing training a deep learning network and specifically focused on bioimages were scarce, mostly Cellpose (14) and StarDist^14^ (7). Finally, 29 projects declared using other tools. A complete list of the individual tools cited in OC 1 can be found in (see Fig. S5a). The categorized answers from OC 2 and 3 are shown in Fig. S5b, including all individual tools from the “Others” category.

**Figure S4.**
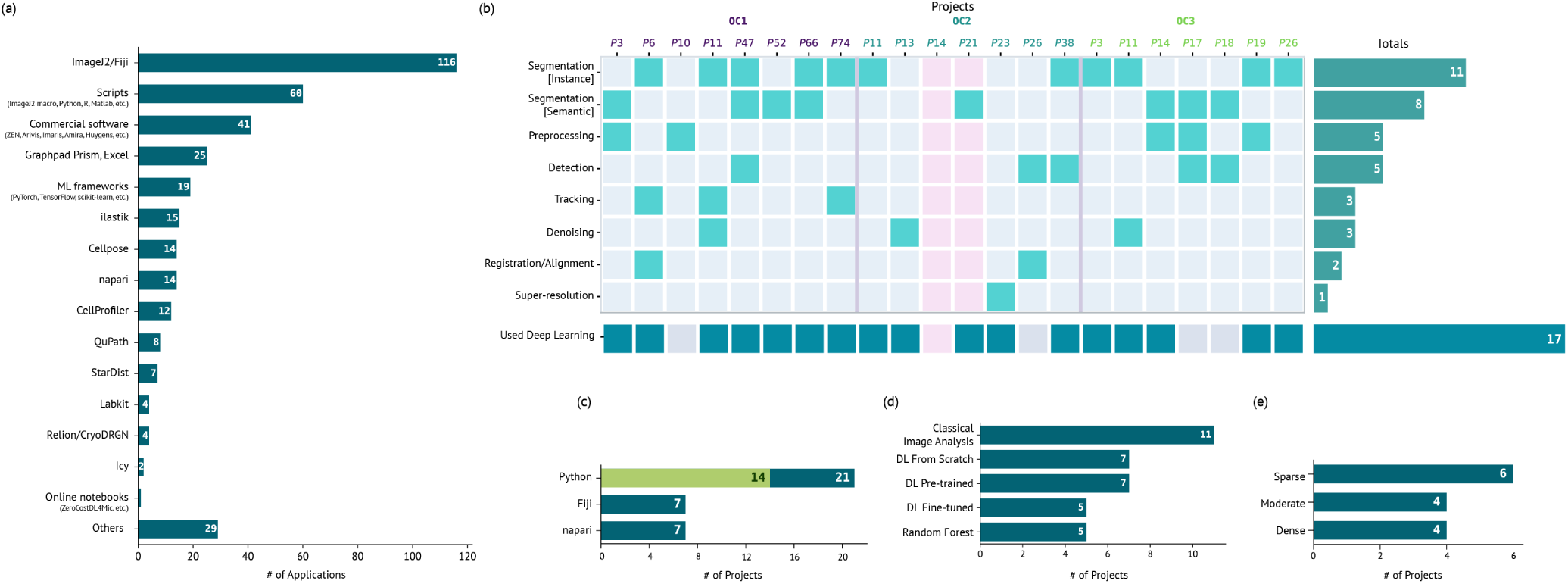
(a) Tools and tool categories declared in the applications as used or tested for the purpose of the project. The “Others” category was available at application time for users to specify additional tools. (b) Discrete analysis steps and their occurrences in every project selected for the three OCs. Red columns indicate failed projects. The right-hand side plot shows the distribution. We indicate in the bottom row whether the project used deep learning. (c) Number of projects using one of three software ecosystems (Python, Fiji, napari). Dark bars indicate to the number of project pipelines partially carried out on the corresponding ecosystem, while green bars denote the number of project pipelines entirely performed in that ecosystem. Project pipelines could make use of multiple software ecosystems. (d) Number of projects using classical image analysis, random forest classification (shallow learning), and various deep-learning flavours (training a model from scratch, using a pre-trained model, and fine-tuning a pre-trained model). Project pipelines could use a mix of these categories. (e) Number of projects requiring various degrees of labeling effort (sparse, moderate, and dense labeling). Sparse labeling corresponds to squiggles typically drawn for training random forests or to point annotations, while moderate labeling to adding a few masks for Cellpose fine-tuning, and dense to complete annotation of a dataset for de-novo training.

#### S2.4 Selected projects

Once projects were selected, expert image analysts were tasked with developing a FAIR pipeline to solve the challenges encountered by the applicants, while ensuring that they could use it in the future. Summary of selected projects can be found in Table S8. In the next sections, we provide an analysis of the pipelines developed for these projects. A description of the resulting pipelines, and links to code and data repositories can be found in Table S9.

#### S2.5 Project Pipelines

The pipelines converged to a limited set of recurring image analysis tasks (see Fig. S4b), dominated by segmentation (19 out of 22 pipelines), pre-processing (5), and detection (5), sometimes combined with denoising (3), tracking (3), or registration/alignment (2). One project focused on super-resolution only. We excluded final quantification as it was the end goal of most projects. Each pipeline relied on different tools and scripts, highlighting the heterogeneity of the analysis goals and the specialization of image analysis tools, as well as preferences of the analysts themselves. The implemented solutions relied on a mix of widely adopted community tools, domain-specific tools and project-specific developments. See Fig. S6 for a breakdown of the various tools and programming/scripting languages used in each image analysis task. All tools used in the OCs are listed in Fig. S7.

All projects used either Python or ImageJ2/Fiji (see Fig. S4c), via custom scripts or tools, or both together in heterogeneous pipelines. Out of the 22 projects, 21 used Python-based tools, of which 14 solutions were Python only (including Python-based tools, light green bars in Fig. S4c). ImageJ2/Fiji was used in 7 projects, but none of the pipelines solely consisted of steps in the ImageJ2/Fiji ecosystem. In on project, the core of the pipeline was carried out in ImageJ2/Fiji, and the Python component was an optional labeling step. napari was also used in 7 projects, 5 times for deep learning related plugins (FeatureForest^15^, nnInteractive^16^, Noise2Void^17^ plugin), one time to train a random forest classifier (APOC^18^), and once for image visualization (OC2-38). A majority of projects required custom scripts to process images or call specialized tools and libraries. The pipelines delivered Jupyter notebooks (13 projects), Python scripts (7 projects), and ImageJ2/Fiji macros (4 projects). Finally, 2 projects delivered containers to the users.

**Figure S5.**
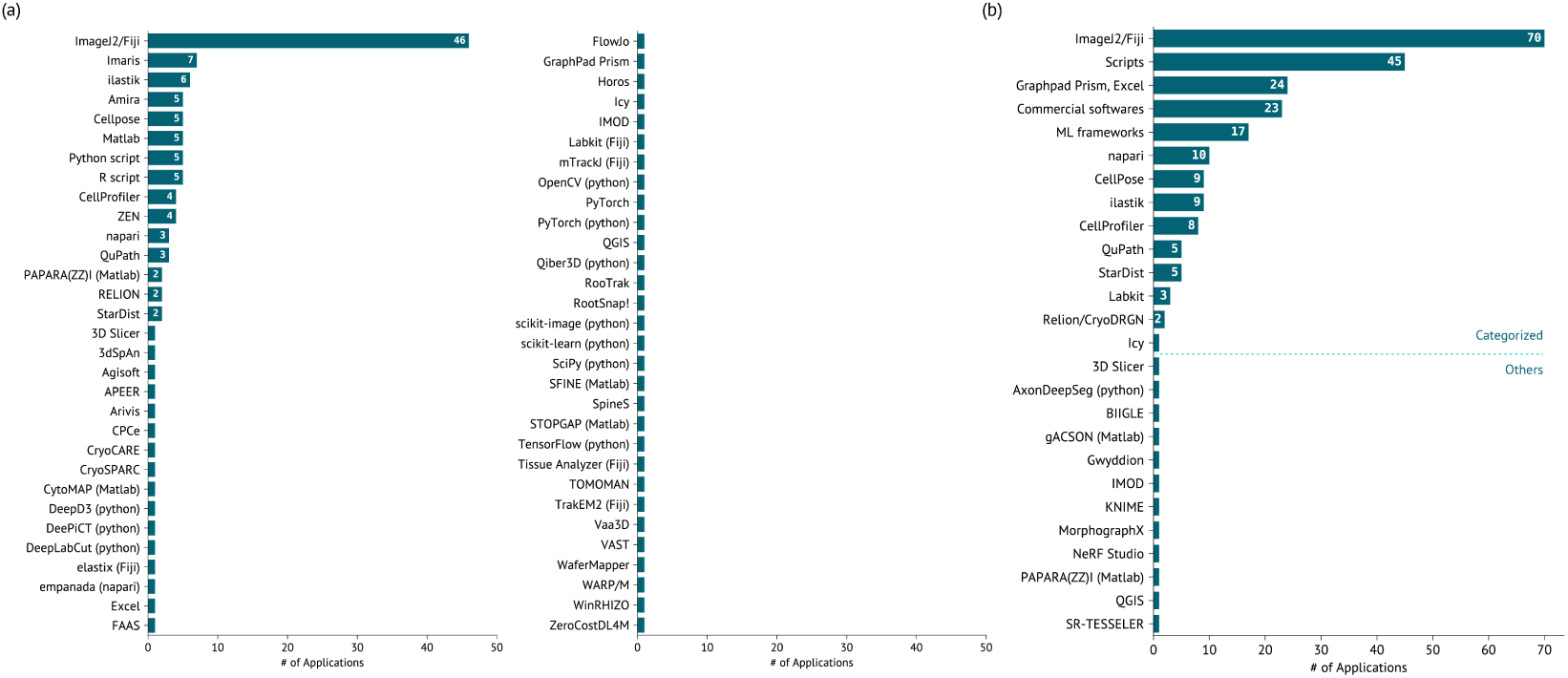
(a) Tools mentioned by users in their OC 1 application as free text. (b) Individual tools and categories of tools selected by users in their OC 2 and 3 application. Additional tools could be mentioned by users as free text (see line separation for “Others”).

**Figure S6.**
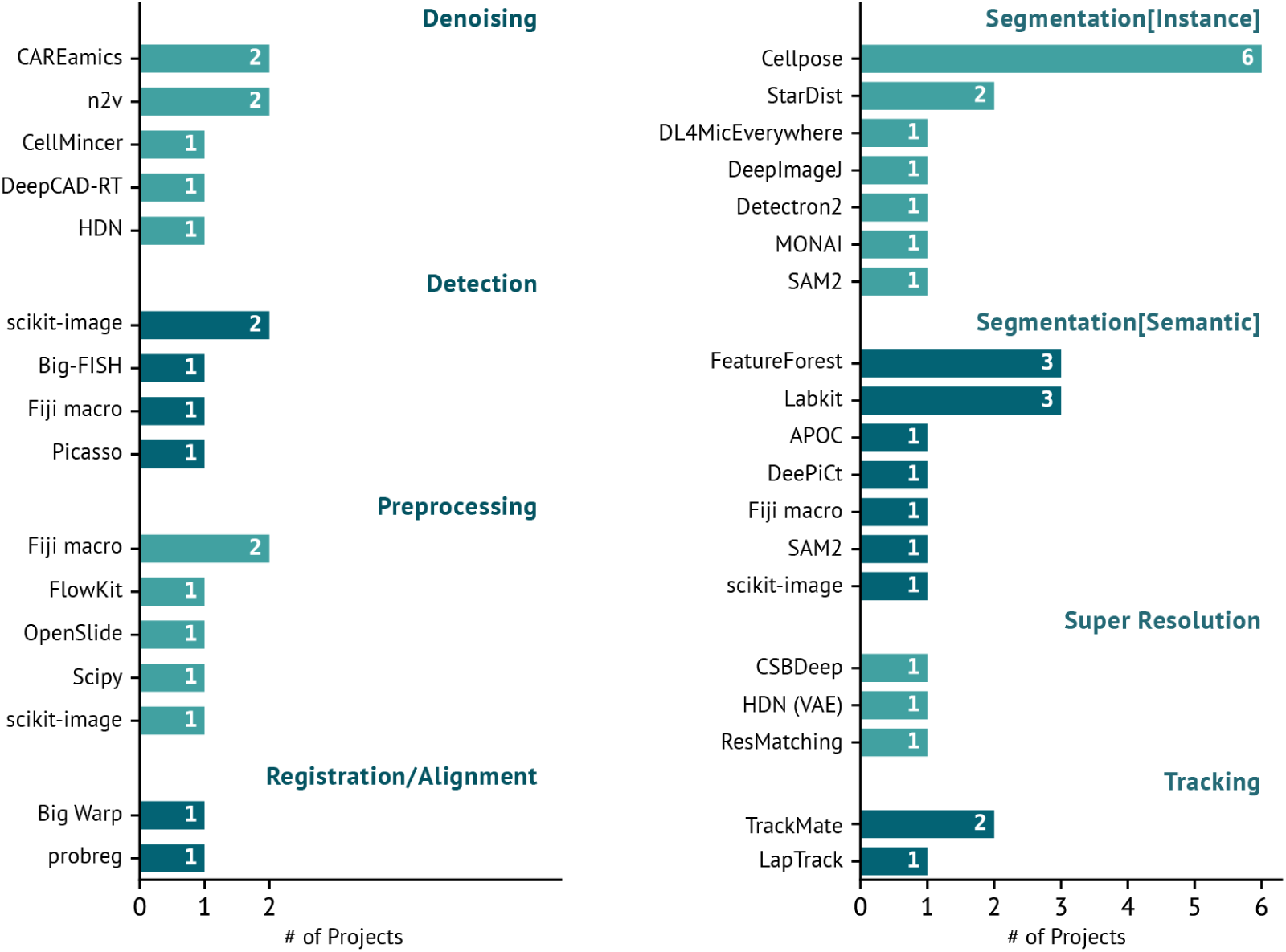
The various tools and libraries used in the project pipelines by analysis steps, to the exclusion of labeling, post-processing and quantification. Project pipelines could make use of several tools in a single analysis step. A project compared denoising algorithms in CAREamics (Noise2Void, Noise2Noise), DeepCAD-RT, CellMincer and HDN. The two instance segmentation steps using StarDist were carried out in DeepImageJ and DL4MicEverywhere. All super-resolution tools were used in a single project.

**Figure S7.**
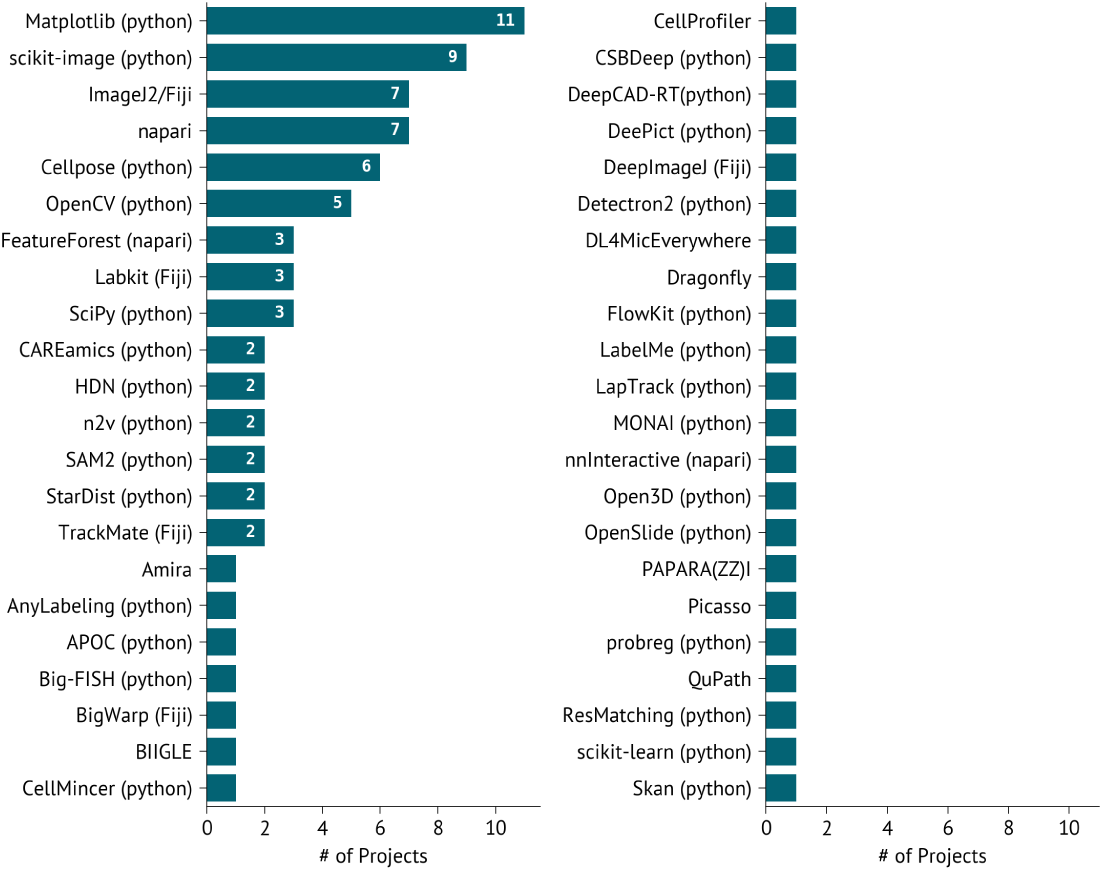
**Summary of tools used in the project pipelines.**

The application of deep learning to bioimages was the focus of the OCs. In the majority of projects, some flavour of deep learning was applied (17 out of 22, see Fig. S4b), while 5 projects also used shallow machine learning (random forests) and 11 projects additionally performed some classical image analysis explicitly (filtering, watershed, morphological operations, etc.). A total of 6 projects employed solely deep learning, without shallow learning or classical image analysis. Random forests pixel classification, without any deep-learning generated features, was mostly carried out in Labkit^12^ (4 projects) and one time with APOC^18^. Classical image analysis was essentially performed in Python (10 projects out of 11), with a mix of scikit-image^19^ (9), OpenCV^20^ (3), SciPy^19^ (2), and Skan^21^ (1), and one time purely in ImageJ2/Fiji. In 3 projects (OC2-26, OC3-17, and OC3-18), the entire pipeline consisted of classical image analysis only, without shallow or deep learning steps.

Some pipeline steps did not involve deep-learning. Pre-processing was performed with ImageJ2/Fiji macros in 2 projects, and with various Python libraries (FlowKit^22^, OpenSlide^23^, SciPy^24^, scikit-image) in three others. Detection was performed using classical image analysis with domain-specific tools (Big-FISH^25^, Picasso^26^), Python scripts using scikit-image (2 projects), or ImageJ2/Fiji macros (1). Registration and alignment were also performed using BigWarp^27^ and probreg^28^. Finally, the last step not performed using deep learning was tracking, although the generation of the instances used for tracking was obtained using deep learning methods. Tracking was present in 3 pipelines and was carried out using TrackMate^29^ (2 projects) or LapTrack^30^ (1).

As expected, deep learning was the approach of choice in denoising, segmentation (instance and semantic) and super-resolution. Among the various pipelines that involved deep learning, pre-trained models were used in 7 projects (Cellpose and SAM2^31^, either directly or via its use within FeatureForest), while 5 projects fine-tuned models (Cellpose) and 7 pipelines trained models entirely from scratch. De-novo trained models were either trained using an existing tool (CAREamics^32^, CellMincer^33^, DeepCAD-RT^34^, DeePiCt^35^, HDN^36^, Noise2Void^17^, StarDist^14^) or custom scripts (3 projects). Denoising was the main focus of project OC2-13 that compared various approaches for the denoising of calcium imaging, leading to the application of CellMincer, DeepCAD-RT, HDN, Noise2Noise^37^and Noise2Void^17^, where the latter two were trained within CAREamics. Two other projects applied denoising using Noise2Void (using the original n2v tool and the more recent CAREamics), demonstrating that it remains the most accessible denoising deep learning-based algorithm. Instance segmentation was dominated by Cellpose (6 out of 11 projects), specifically Cellpose2^8^. StarDist was used twice, once from within DeepImageJ^38^ via CellProfiler^10^ and the other using DL4MicEverywhere^39^. The other instance segmentation approaches were a custom training of Detectron2, of a U-Net^40^ from Monai^41^, and an application of SAM2^31^. Semantic segmentation with deep learning was either carried out using random forests trained from the embeddings of SAM2 (FeatureForest^15^, 3 projects), or a de-novo trained model (DeePiCt, 1 project). Other semantic segmentation steps were done using random forests directly (Labkit^12^ in 3 projects, APOC^18^ jointly in one of these projects). Finally, the super-resolution project compared several methods: CARE^42^ (using the CSBDeep library), ResMatching^43^ and the variational auto-encoder (VAE) from HDN.

While the deep learning methods used in the OC pipelines often differ from each other by their purpose and features, the majority of the methods were UNet-based (13 out of 17, see Fig. S8), with 4 visual transformers-based approaches (SAM2-derived) and two other architectures (Mask R-CNN, variational auto-encoder).

**Figure S8.**
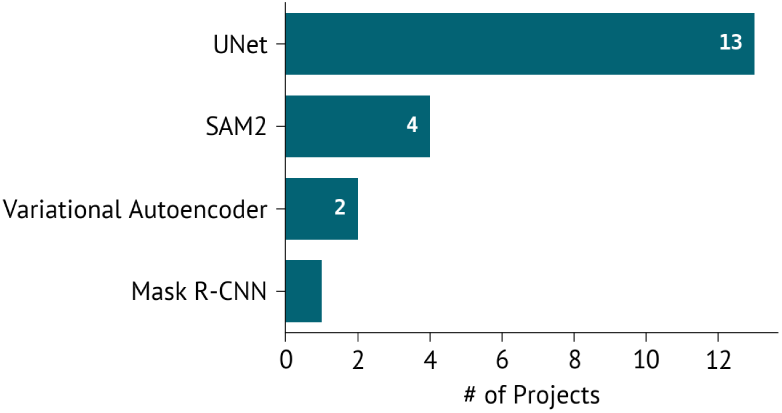
Deep learning architectures or framework used in the OC pipelines. Note that in one project, SAM2 encoder was used both in FeatureForest and in a hybrid SAM2+UNet model.

As a consequence of the prevalence of segmentation in the analysis steps, 13 of the 22 projects required annotations (see Fig. S4e). Since gold-standard annotations were scarcely available (see next section), methods requiring a sparse or moderate amount of annotations were often adopted (6 out of 22). That was specifically the random forests-based pipelines, Labkit and FeatureForest, which required sparse squiggles-like annotations, and the point labels used for SAM2. Cellpose fine-tuning only necessitated a few additional masks, corresponding to a moderate labeling effort (4 projects). De-novo training of a deep learning network for segmentation purposes, on the other hand, required dense annotations (5 projects). A single project had enough gold standard annotation at the start of the project to train a deep neural network (OC3-14), albeit only for one class out of the four originally expected to be segmented. Annotations were either performed in the segmentation tools directly (Labkit, APOC, FeatureForest, Cellpose), in software with annotation tools (napari^9^, BIIGLE^44^, Amira^45^, Dragonfly^46^, LabelMe^47^, AnyLabeling^48^), or with deep learning tools (nnInteractive^16^). Of the aforementioned tools, Labkit is a ImageJ2/Fiji plugin, while APOC, FeatureForest, and nnInteractive are napari plugins. In two cases, user annotations had already been generated in PAPARA(ZZ)I^49^ and QuPath^11^, and these tools were used to export the annotations.

Finally, image visualization was often performed with ImageJ2/Fiji (5 projects) or napari (6). Many Python scripts and notebooks used Matplotlib ^50^ to visualize results (11 projects), and one notebook employed Bokeh^51^. Domain-specific tools were also used to check results, with the example of BIIGLE^44^ (1 project), CellProfiler^10^ and Picasso^26^ (1).

#### S2.6 Project readiness and outcome analysis

Only 8 out of 22 selected projects were fully ready at the start of the support period, with complete, usable data and sufficient annotations available to start with. The remaining projects required some form of intervention before or during execution: additional data acquisition, re-imaging, or further ground-truth annotation.

Data shareability was assessed at the application stage across all 151 applicants: 93 indicated their data was fully shareable, 53 partially shareable, and only 5 not shareable (see Fig S9). In OC 2 and 3, the option to declare the data as non-shareable was not available as it was a mandatory outcome of the OC projects.

**Figure S9.**
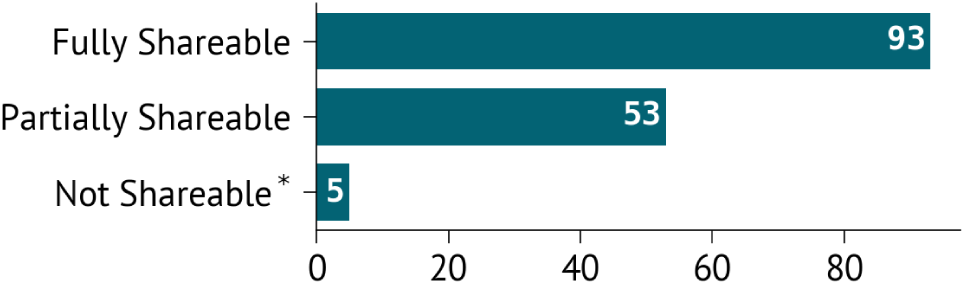
**Portion of their data that is shareable as declared by the applicants. “Not Shareable” was only available as an answer in OC 1.**

Data transfer to the image analysis team was a recurring bottleneck, with 3 out of 22 projects experiencing significant delays. Delays arose both from applicant-side constraints, including slow uploads and late data submission, and from technical limitations, such as the absence of a unified data-transfer solution across institutions. Across 22 projects, 14 different data-transfer methods were used, ranging from commercial cloud services to institutional platforms and one-time physical hard drives sent via postal services (see S10), highlighting the fragmentation of current data-sharing practices.

**Figure S10.**
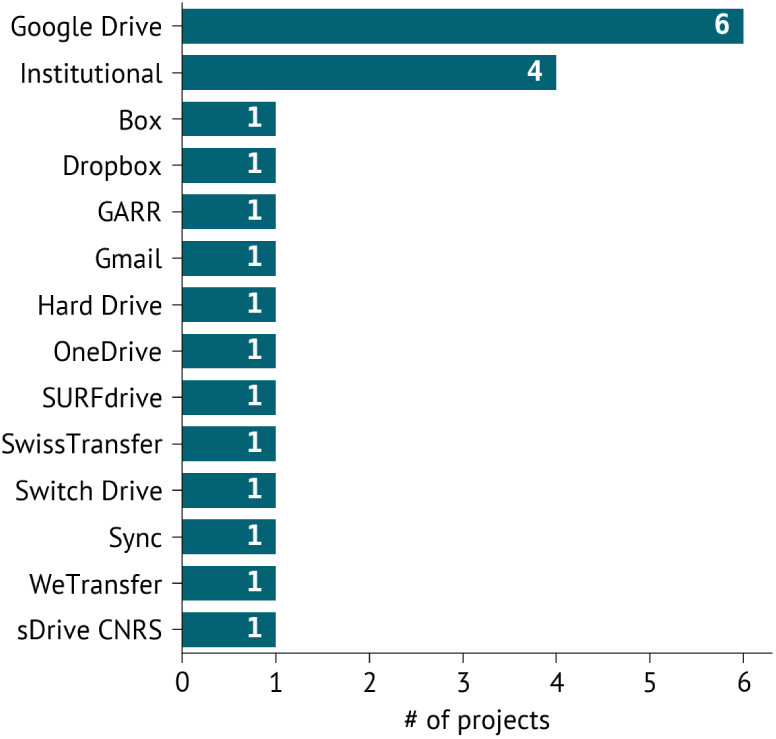
Data transfer methods used during project support as declared by the image analysts. Institutional refers to any service provided by the home institution of the applicants or the analysts. “Hard Drive” refers to a physical hard drive sent by the post.

All projects were assessed at the end of the project by the support team experts for data readiness for deep learning, taking into account annotation quantity, quality, and coverage. Only 2 out of 22 projects were considered to have sufficient ground-truth data for deep learning. Among the others, 3 projects had no ground-truth annotations available at the start of the support period, and 3 further projects required substantially more annotation than originally provided during project runtime to reach the intended analysis objectives. In 4 out of 22 projects, data quality was lower than initially expected, causing delays and requiring corrective steps due to corruption, saturation, imaging noise, or other acquisition issues. All readiness and quality assessments reported here were filled out by the experts directly involved in supporting each project.

A small number of projects (2 out of 22) did not achieve the initially defined objectives by the end of the support period. In these cases, outcomes were limited by factors such as insufficient data readiness, overly ambitious or evolving project scope, or technical issues encountered during execution.

All projects pipelines, except for two failed projects (OC2-14 and OC2-21), one under publication embargo (OC2-23), and one currently in its final steps (OC3-26), were uploaded to GitHub with scripts, notebooks, or detailed descriptions on how to reproduce the pipeline. The high-level description and results of the project under publication embargo are available in place of the complete pipeline. All GitHub repositories were also archived on Zenodo. For 13 of the projects, data was uploaded to the BioImage Archive, while 4 were uploaded to Zenodo (see Fig. S12). Illustrating the difficulty in transferring data, track metadata and incentivize users in publicly sharing it, we observed significant delays in sharing the data publicly (see Fig. S11). 17 out of 22 projects led to publicly released data, while one project is still awaiting data upload. 4 projects had no useful data to upload: 2 failed projects, 1 software-only project for which data was not applicable (OC3-3), and 1 case where contact with the applicant was lost after the project had ended (OC1-47). To quantify the delays in releasing public data, we used two indicators: the official end of support project and the last update to the public pipeline. Indeed, project support may have been extended beyond the official support time frame due to multiple factors: incurred delays in transferring or acquiring data, final steps requiring more polishing, issues in the pipeline surfacing after official support has ended, etc.. Data was publicly available within a month of the end of the official support for 1 project only (OC3-19, see Fig S11a), while the data for another 11 projects (out of 17) took 6 months or more to be uploaded, of which 3 took a year or more. The average time from the end of official support to public data release was 8.1 months. The last changes to the pipelines were consistently done after the end of the official release. Consequently, the distribution of estimated time between the last pipeline modification and data release leads to shorter time frames (see Fig S11b). Nonetheless, the average time to public data release remains sizable in that case: 4.4 months. Only 2 projects saw data being released at the same time as the final touches to the public pipeline (OC1-6 and OC1-10). 9 datasets were uploaded online 3 months or more after the pipeline was finalized, of which 2 took 7 months and another 2 more than a year. Note that 2 projects, OC2-23 and OC3-26, do not have yet a public pipeline as one is currently under embargo for publication (OC2-23) and the other one is not yet finalized (OC3-26).

Finally, trained models were uploaded to the BioImage Model Zoo for 5 projects (OC1-6 OC1-11, OC1-66, OC1-74, and OC3-11), while 2 projects’ models were uploaded to Zenodo (OC2-13, OC3-19), and 1 additional random forest model was deposited alongside its dataset on the BioImage Archive (OC1-3). Pre-trained models that were not fine-tuned were not uploaded to the BioImage Model Zoo. Links to the pipelines, data and models can be found in Table S9.

**Figure S11.**
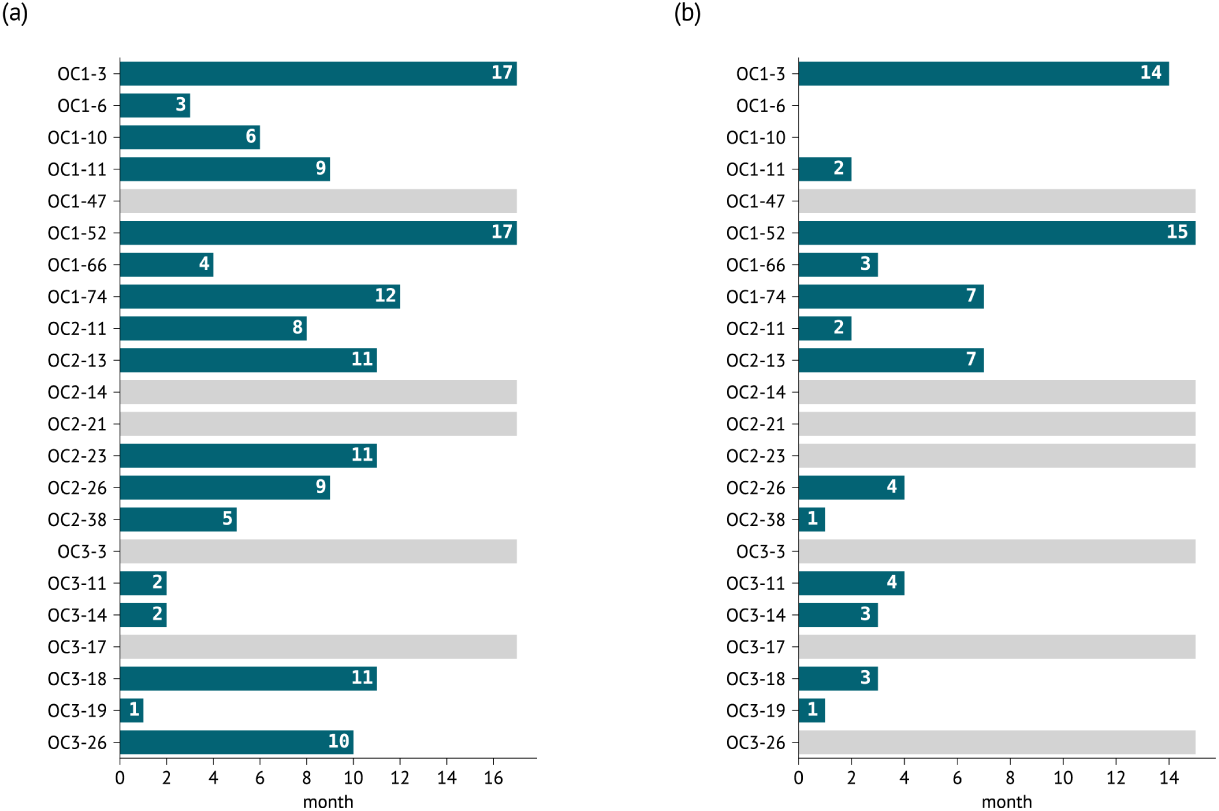
(a) Time in months between end of official support and public data release for all projects. Grayed-out projects do not have publicly released data, due to loss of contact (OC1-47), project failure (OC2-14 and OC2-21), not having data (OC3-3), and data in the process of being uploaded at the time of writing (OC3-17). (b) Time in months between last update of the public pipeline and public data release for all projects. Grayed-out projects do not have publicly released data (see (a)) or pipeline. OC1-6 and OC1-10 coincidentally released data and carried out the last update to the pipeline, while OC2-23 (embargo) and OC-26 (ongoing) have released data but no public pipeline at the time of writing.

**Figure S12.**
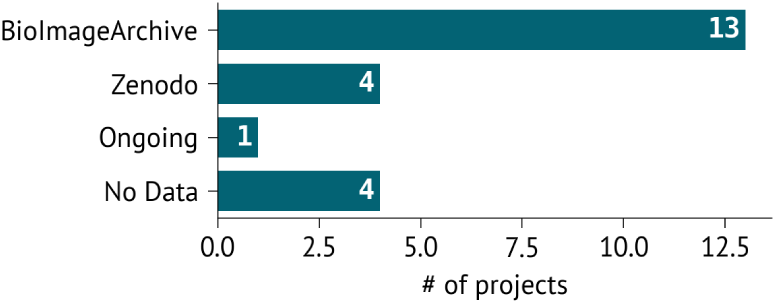
Repository used for openly released data. “Ongoing” refers to selected project whose data is not yet released at the time of writing. “No Data” corresponds to the two failed projects, one lost contact, and one project whose aim was to write a software plugin.

#### S2.7 Methods

All the application stats were extracted from the application forms filled by applicants across the three open calls. All the responses were concatenated together in an Excel sheet and mined directly or otherwise curated. Some categories resulted in lists of entries (e.g., data formats, imaging methods, image analysis tools), which were then individually split. We then manually curated the answers to standardize the terms (e.g. “Lighsheet” vs “light-sheet”). Nationalities (only OC 2 and 3) and countries of home institution (Fig. S2a and S3) were compiled directly from user answers. Since the scientific domains (Fig. S2c) where free text, users could enter answers that were not strictly speaking scientific domains (e.g. confocal imaging). In a second-pass curation, we removed any such entry. The list of available data formats was a detailed list in OC 1 (specific formats and a free text field), while it corresponded to the general categories shown in Fig. S2e in OC 2 and 3. We therefore automatically allocated the answers of OC 1 into the categories of OC 2 and 3. The type of home organization (Fig. S2b) was manually investigated based on web searches.

Imaging methods (Fig. S2d) were not a field in the application, and were manually derived from specific mentions in the project description field, data description field, project summary presentation, and investigation of uploaded data. Wherever ambiguity could not be lifted, the applications were categorized as “Others”. CLEM application was split into “Widefield” and electron microscopy “Others” (transmission electron microscopy). One successful application mentioned “SHG” among other imaging methods but the project did not make use of that particular data, it was therefore excluded from the statistics. While there are overlaps between categories (e.g., SMLM and Widefield, Fib-SEM and vEM) or techniques that are strictly imaging methods (e.g., Expansion), we reported the denomination used by the applicants. TEM and SEM added ambiguity in the classification of EM methods and where aggregated into “Others” as a result. Natural images (e.g. underwater photography, seed pictures) were aggregated into a single “Photography” category. Scanography refers to images generated by using a scanner.

Tools declared by the applicants required manual curation and web search to confirm the existence of the tools/software/library and their correct spelling. In OC 1, tools were declared as free text. Based on the statistics from OC 1, we introduced individual and categories of tools in the application form OC 2 and 3. In order to plot Fig. S4a, we categorized the answers from OC 1 using the same categories as in OC 2 and 3.

To compile the statistics of the selected projects (Fig. S4b, c, d and e), we investigated each GitHub repository to extract analysis tasks, programming language, tools, library and software usage, labeling effort, deep learning usage, training mode (pre-trained model, fine-tuning, de-novo training).

Project outcomes and data readiness were assessed by the support team member assigned to each project and recorded at the end of the support period. Each assessment included explicitly structured fields: a readiness score on a 1–5 scale, the support team’s judgment of whether sufficient ground-truth data was available for deep learning, the self-reported ground-truth status provided by the applicant at project initiation, the data transfer method used, and the overall project outcome. In addition, each assessor provided a free-text comment describing any issues encountered during the project. The statistics on data transfer delays, re-imaging events, annotation insufficiency, and data quality problems reported in this section were derived from systematic analysis of these free-text comments.

To estimate the delay between the end of project and public data release for the 16 projects that did release data, we used two different end of project dates. The first one is the official end of project support time frame: December 31*^st^* 2023 (OC 1), October 31*^st^* 2024 (OC 2), and August 31*^st^* (OC 3). However, projects may have incurred delays and actual project support be extended in time. We therefore added an estimated end of project support using the latest git commit modifying the public pipeline. Public data release was readily available from BioImage Archive and Zenodo releases. The two projects with no public pipelines were excluded from the average calculation in the case of the end of support date estimated from the pipeline commits.

### S3 Public Challenges

#### Problem selection

During the Open Calls we observed that biological image analysis pipelines typically involved preprocessing, often denoising, followed by segmentation and downstream analysis. While segmentation remains a key step, state-of-the-art tools such as Cellpose^7^ and StarDist^14^ already perform robustly on many common biological imaging tasks. Downstream analysis, by contrast, tends to be highly project-specific, making it difficult to formulate it as a general-purpose challenge.

Image restoration, particularly denoising, stood out as a more broadly applicable and impactful task. Improvements in denoising directly influence the performance of subsequent steps, such as segmentation and quantification, and enable the effective use of low-signal or low-quality data, which is common in biological imaging. We therefore identified a clear need for improved denoising methods and for systematic benchmarking of existing approaches, and more broadly, for objective comparison of methods under controlled, reproducible conditions, which is a prerequisite for understanding the true state of the field.

Depending on the data at hand, image denoising may require fundamentally different approaches. We therefore designed three challenges to cover complementary denoising settings, each targeting a distinct data regime.

#### Hosting platform selection

Hosting a deep learning challenge requires robust infrastructure for dataset management, solution submission, automated evaluation, and public leaderboards. Several established platforms (Kaggle, Grand Challenge) already provide these capabilities. In addition to technical considerations, established platforms offer built-in visibility and access to an engaged user base, which is critical for reaching the broader machine learning community.

After a comparative evaluation, we selected Grand Challenge, a European platform specializing in medical imaging challenges. Although no existing platform focuses exclusively on biological imaging, Grand Challenge offered the closest alignment with our requirements. Its emphasis on medical imaging, support for reproducible Docker-based submissions and compliance with European data regulations made it a strong fit.

#### Organization process

Each challenge began with the definition of a concrete and achievable task, which was formulated to be understandable to participants without prior experience in bioimage analysis. Appropriate datasets were then selected based on multiple criteria, including sufficient size for training deep learning models, alignment with the challenge objective, coverage of different imaging modalities, and representation of diverse biological sample types.

For each challenge, we prepared comprehensive documentation describing the scientific motivation, task definition, dataset characteristics, relevant background literature, and detailed instructions for accessing the data and submitting solutions via the hosting platform. To lower the barrier to participation, we also provided starter code and clear instructions for packaging submissions using Docker. Throughout the challenge periods, ongoing technical support was offered, including assistance with submission issues, clarification of task requirements, and guidance related to the challenge infrastructure.

Top-performing participants were required to release their training code publicly as a condition of recognition and were invited as co-authors of this summary paper. Participants who did not publish their code or who could not be reached for follow-up are acknowledged instead.

#### Challenges organized

he three challenges were unsupervised denoising (PC 1), supervised denoising (PC 2), and domain-specific denoising for calcium imaging (PC 3). In PC1, the data consisted of noisy data only, while PC2 focused on methods trained on noisy-clean image pairs. PC3 required methods to preserve temporal signal structure in addition to spatial image quality, and was inspired by an Open Call project (OC2-13).

Each challenge included four leaderboards corresponding to different datasets or task variants (full details are provided in Table S1). Datasets were selected from openly available sources with ground truth, or, where new data were introduced, training data were made publicly available on Zenodo. To ensure fair evaluation, test sets for leaderboards with ground-truth labels were privately hosted on the platform and not released to participants.

**Table S1.** Datasets used in the Public Challenges.

| Challenge | Leaderboard | Dataset | Imaging modality | Dim. | Training data |
| --- | --- | --- | --- | --- | --- |
| Unsupervised Microscopy Denoising Challenge | Hagen et al.: Structured noise | Hagen et al.<sup>52</sup> [data] | Confocal microscopy | 2D | 79 images |
|  | SUPPORT: Structured noise | SUPPORT<sup>53</sup> [data] | Confocal microscopy | 3D | 1 volume (1001 z-slices) |
|  | W2S: Unstructured noise | W2S<sup>54</sup> [data] | Widefield microscopy | 2D | 120 images |
|  | JUMP: Unstructured noise | JUMP<sup>55</sup> [data] | Fluorescence microscopy | 2D | 517 images |
| Supervised Microscopy Denoising Challenge | Planaria: 3D denoising | Planaria<sup>42,56</sup> [data] | Confocal microscopy | 3D | 17,945 paired 3D crops |
|  | Tribolium: 3D denoising | Tribolium<sup>42,56</sup> [data] | Multiphoton laser-scanning | 3D | 15,500 paired 3D crops |
|  | Nuclei: 2D denoising | Nuclei<sup>57,58</sup> [data] | Widefield/spinning disk confocal | 2D | 2,477 image pairs |
|  | FMD: 2D denoising | FMD<sup>59</sup> [data] | Confocal/two-photon/wide-field | 2D | 63,669 image pairs |
| Calcium Imaging Denoising Challenge | All leaderboards | AI4Life-CIDC25 [train data, val data] | Synthetic calcium imaging | 3D | 5 time-series volumes |

All submissions were evaluated automatically: PC 1 and PC 2 were scored using scale-invariant PSNR, while PC 3 employed a spatiotemporal PSNR metric to account for the temporal component of the signal (see methods).

### S4 Public Challenges outcomes

#### S4.1 Challenge statistics

In total, the three challenges attracted 225 registered participants from 41 countries and resulted in 145 successful submissions (with 289 total submission attempts). Participation was geographically diverse, with the largest contingents from China (44), India (28), the United States (27), the United Kingdom (15), and Italy (12), alongside participants from 36 additional countries. These statistics are represented in Table S2.

**Table S2.** Countries by registrant count. Other countries: Germany (5), Spain (5), Switzerland (5), Bangladesh (4), Iran (4), Pakistan (4), Belgium (3), Canada (3), Czechia (3), Sweden (3), Taiwan (3), Brazil (2), Colombia (2), Ecuador (2), Israel (2), Armenia (1), Australia (1), Denmark (1), Egypt (1), Greece (1), Indonesia (1), Ireland (1), Malaysia (1), Morocco (1), Nepal (1), Nigeria (1), Portugal (1), Romania (1), Slovakia (1), Sri Lanka (1), and Tanzania (1).

| Country | Registrants |
| --- | --- |
| China | 44 |
| India | 28 |
| United States of America | 27 |
| United Kingdom | 15 |
| Italy | 12 |
| Hong Kong | 7 |
| South Korea | 7 |
| France | 6 |
| Türkiye | 6 |
| Vietnam | 5 |
| Other | 66 |

Engagement varied across challenges. The Unsupervised Microscopy Denoising Challenge (PC 1) drew the largest audience, with 148 registrants and 68 successful submissions. The Supervised challenge (PC 2) attracted far fewer registrants (19) but produced 34 successful submissions, the highest submission-to-registrant ratio of the three, suggesting a highly engaged participant base. The Calcium Imaging Denoising Challenge (PC 3) attracted 58 registrants and had 43 successful submissions out of 72 total attempts, which we attribute to the increased task difficulty: calcium imaging requires handling volumetric data under complex noise conditions, likely demanding substantially more effort from participants. Reflecting this, PC 3 ran for considerably longer than the other two challenges, with the submission window extended multiple times in response to slow initial uptake. Overall, the number of successful submissions remained modest relative to established challenges in computer vision or medical imaging. The statistics are presented in Figure S13. Full submission counts per challenge and per leaderboard are also provided in Table S3 and S4.

**Table S3.** Registrants and submissions per challenge.

| Challenge | Period | Registrants | Total Submissions | Successful Submissions |
| --- | --- | --- | --- | --- |
| Unsupervised Microscopy Denoising Challenge | Jun-Jul 2024 | 148 | 161 | 68 |
| Supervised Microscopy Denoising Challenge | Jun-Aug 2025 | 19 | 56 | 34 |
| Calcium Imaging Denoising Challenge | Jun 2025-May 2026 | 58 | 72 | 43 |
| **Total** |  | **225** | **289** | **145** |

Within each challenge, 2D denoising tasks consistently attracted more submissions than their 3D counterparts. In the Supervised challenge (PC 2), for example, the Nuclei (2D) leaderboard received 14 successful submissions versus 5 for Planaria (3D).

**Figure S13.**
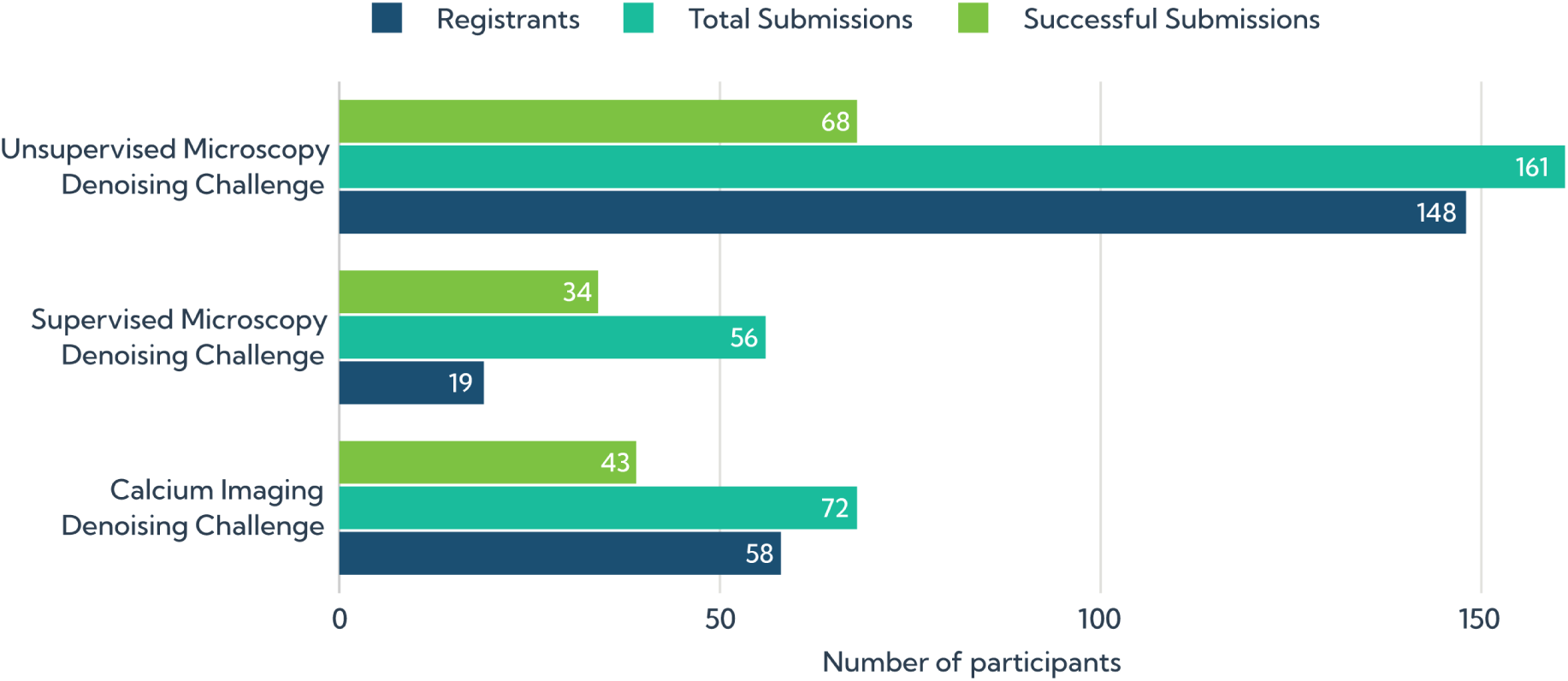
Participation statistics across the three AI4Life denoising challenges. Each Challenge is broken down in three categories: number of registrants, total number of submissions, and number of successful submissions.

#### S4.2 Submitted solutions

Across all three challenges, submissions were dominated by a small number of established method families. Noise2Void^17,32^ and its variants were the most widely adopted approach, appearing across all three challenges and encompassing a range of extensions including probabilistic, structured, and 3D variants. U-Net-based architectures, including a ResUNet^60^ and a method based on CARE^42^, appeared exclusively in the supervised challenge (PC 2). Noise2Fast^61^ and Noise2Noise^37^ variants each contributed two submissions. Noise2Noise-based approaches appeared in the unsupervised and calcium imaging settings. A small number of submissions used more modern and specialized architectures, including COSDD^62^, HDN^36^, a transformer-based model, and a classical method (a correlation-weighted spatial filter).

The algorithm types used are mentioned in Table S5.

Leaderboard results revealed several notable patterns. In the unsupervised challenge, COSDD (University of Birmingham) placed at or near the top across all four leaderboards, consistently outperforming the dominant Noise2Void-based submissions. In the supervised challenge, NAFNet-GAN^63^ (Universidad Carlos III de Madrid) ranked first on both 3D and one of the 2D leaderboards, while placing third in the last leaderboard. Notably, a small number of participants placed consistently across multiple leaderboards within the same challenge, suggesting that general-purpose implementations can perform competitively across diverse datasets. The top 3 participants are shown in Table S6.

Many leaderboard entries corresponded to iterative variants of the same underlying approach, reflecting parameter tuning and dataset-specific adaptations rather than fundamentally distinct methods. Overall, the submitted solutions provide a representative snapshot of current denoising practices across different supervision regimes and data modalities.

Among the top-3 participants across all three challenges, 13 open-source repositories were published containing training code for reproducibility of the results. All are archived on Zenodo to ensure long-term availability, and links are compiled in Table S7.

#### S4.3 Methods

##### Submission and evaluation

Participants packaged their trained models and inference code as Docker containers. To lower the barrier to entry, we provided a template container with example code and detailed submission instructions for each challenge. Submissions were executed on the Grand Challenge platform against privately held test sets not released to participants. Predicted outputs were then evaluated using a dedicated metrics container developed by the organizers.

##### Metrics

PC 1 and PC 2 were scored using scale-invariant PSNR (SI-PSNR), a variant of the standard peak signal-to-noise ratio that is invariant to the scale of the predicted signal.

PC 3 was ranked using a spatiotemporal SNR metric (stSNR), combining a spatial SNR computed per frame with a temporal SNR computed per spatial location across time. This composite metric captures both image quality and the preservation of temporal signal dynamics, which is critical for calcium imaging where the temporal trace carries the biological signal of interest.

Full details and metric formulas are available on the respective challenge pages.

**Table S4.**
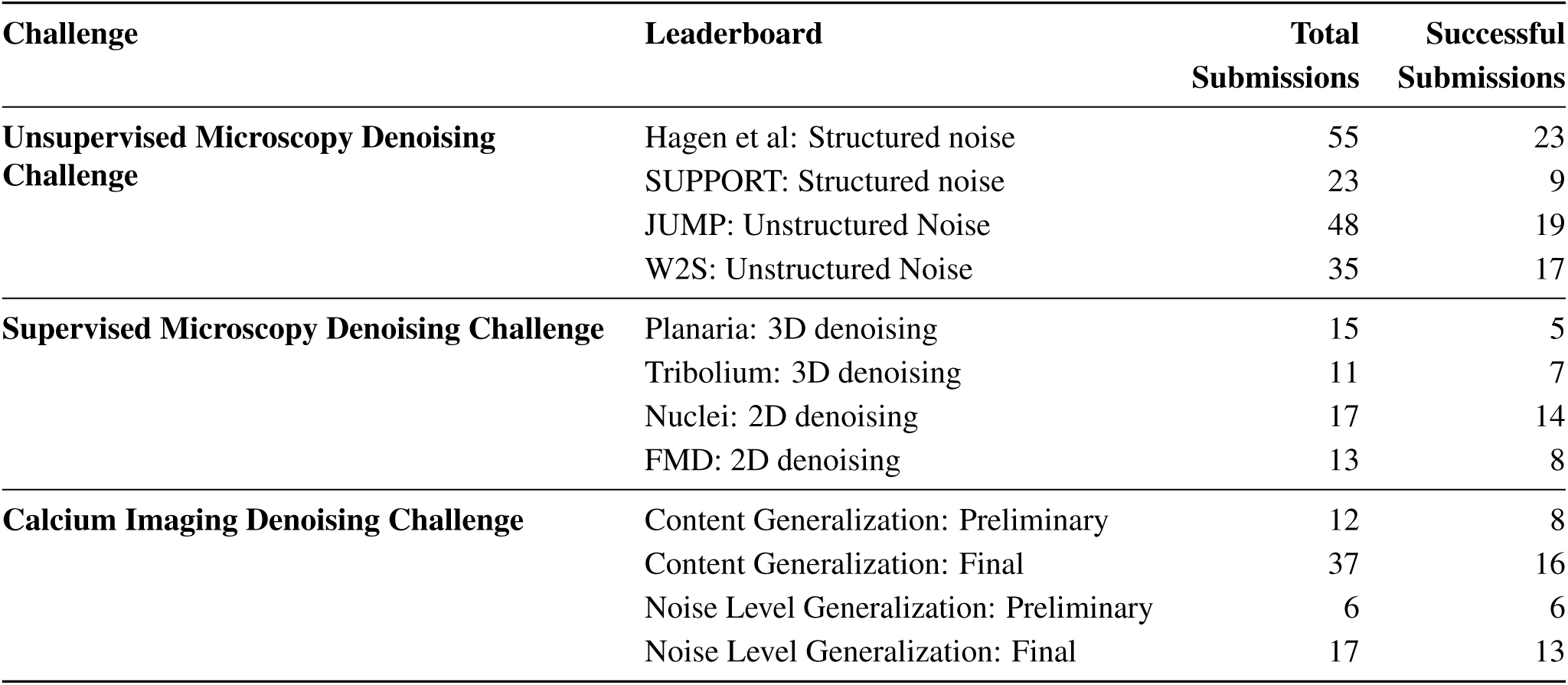
Challenge leaderboard submissions.

**Table S5.**
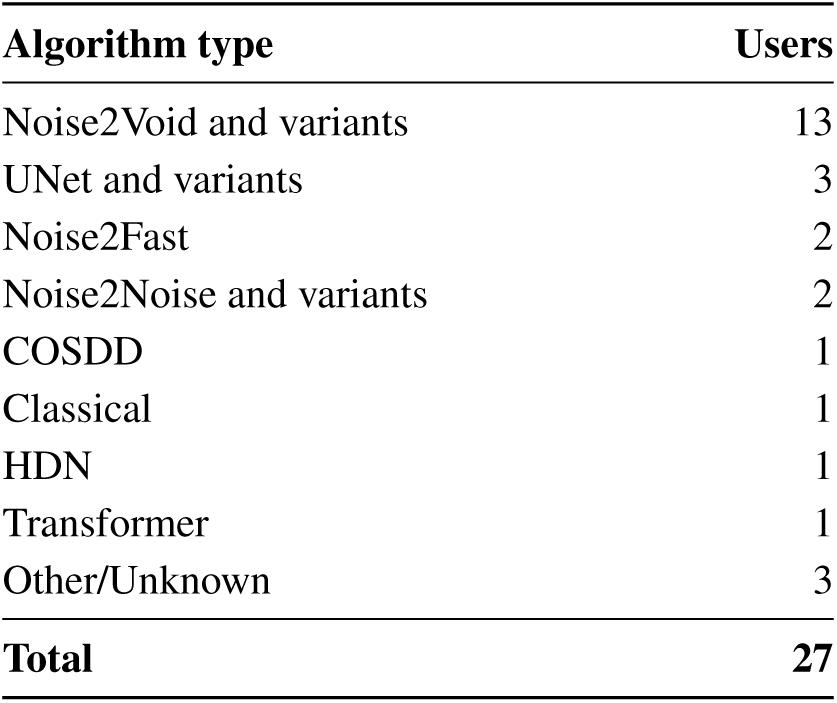
Algorithm types used across challenges.

| Algorithm type | Users |
| --- | --- |
| Noise2Void and variants | 13 |
| UNet and variants | 3 |
| Noise2Fast | 2 |
| Noise2Noise and variants | 2 |
| COSDD | 1 |
| Classical | 1 |
| HDN | 1 |
| Transformer | 1 |
| Other/Unknown | 3 |
| **Total** | **27** |

##### Challenge statistics

Raw submission and participation data were downloaded directly from grand-challenge.org. Country and institutional affiliation were taken from user profile information provided on the platform. A submission was counted as successful if both the inference container and the metrics evaluation container completed without error and produced an output in the expected format.

Algorithm types were manually inferred from the metadata provided by participants at submission time (algorithm name and comment fields) and, where available, from the linked code repositories. Submissions were assigned to algorithm families (e.g., Noise2Void and variants, U-Net and variants) based on this information. For the algorithm family summary Table S5, submissions were de-duplicated such that each unique combination of challenge, participant, and algorithm family was counted once.

**Table S6.**
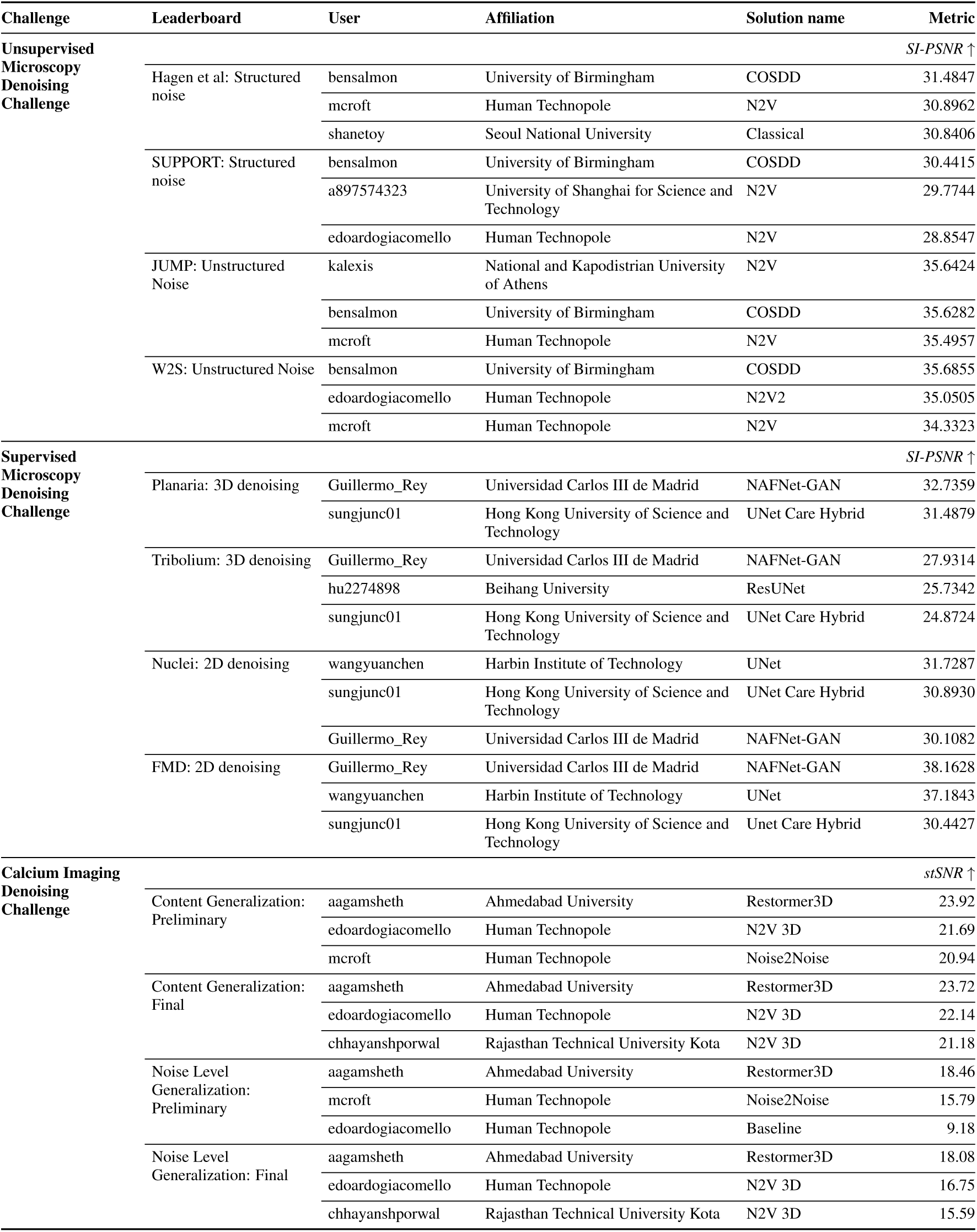
Top 3 participants per leaderboard, showing only the best-scoring submission per participant. For Challenge 3, the challenge administrator (edoardogiacomello) placed in the top 3 on all leaderboards; his results are included as a baseline reference, allowing readers to contextualise participant performance relative to an expert submission.

**Table S7.** Code repositories for all top-3 participants for every leaderboard, grouped by challenge. “—” indicates no code was submitted.

| Challenge | Username | Affiliation | Solution |
| --- | --- | --- | --- |
| **Unsupervised**<br>**Microscopy**<br>**Denoising** | bensalmon | University of Birmingham | GitHub |
|  | mcroft | Human Technopole | GitHub |
|  | shanetoy | Seoul National University | GitHub |
|  | a897574323 | Univ. of Shanghai for Sci. & Tech. | GitHub |
|  | edoardogiacomello | Human Technopole | GitHub |
|  | kalexis | Nat. & Kapodistrian Univ. Athens | — |
| **Supervised**<br>**Microscopy**<br>**Denoising** | guillermo_rey | Univ. Carlos III de Madrid | GitHub |
|  | sungjunc01 | Hong Kong Univ. of Sci. & Tech. | GitHub |
|  | hu2274898 | Beihang University | GitHub |
|  | wangyuanchen | Harbin Institute of Technology | — |
| **Calcium**<br>**Imaging**<br>**Denoising** | aagamsheth | Ahmedabad University | GitHub |
|  | edoardogiacomello | Human Technopole | GitHub |
|  | mcroft | Human Technopole | GitHub |
|  | chhayanshporwal | Rajasthan Technical Univ. Kota | GitHub |
|  | guillermo_rey | Univ. Carlos III de Madrid | GitHub |

The top-3 Table S6 reports the single best-scoring submission per participant per leaderboard. Where a challenge administrator placed in the top three (PC 3), their result is included as a reference baseline and noted as such in the table caption.

## A1 Appendix: OC3 application form

The form had five short sections in which we tried to understand the needs of the potential project:

- Applicant personal information
- General information about the project
- The existing image data
- The current image analysis workflow (if any)
- Questions about potential impact and dissemination

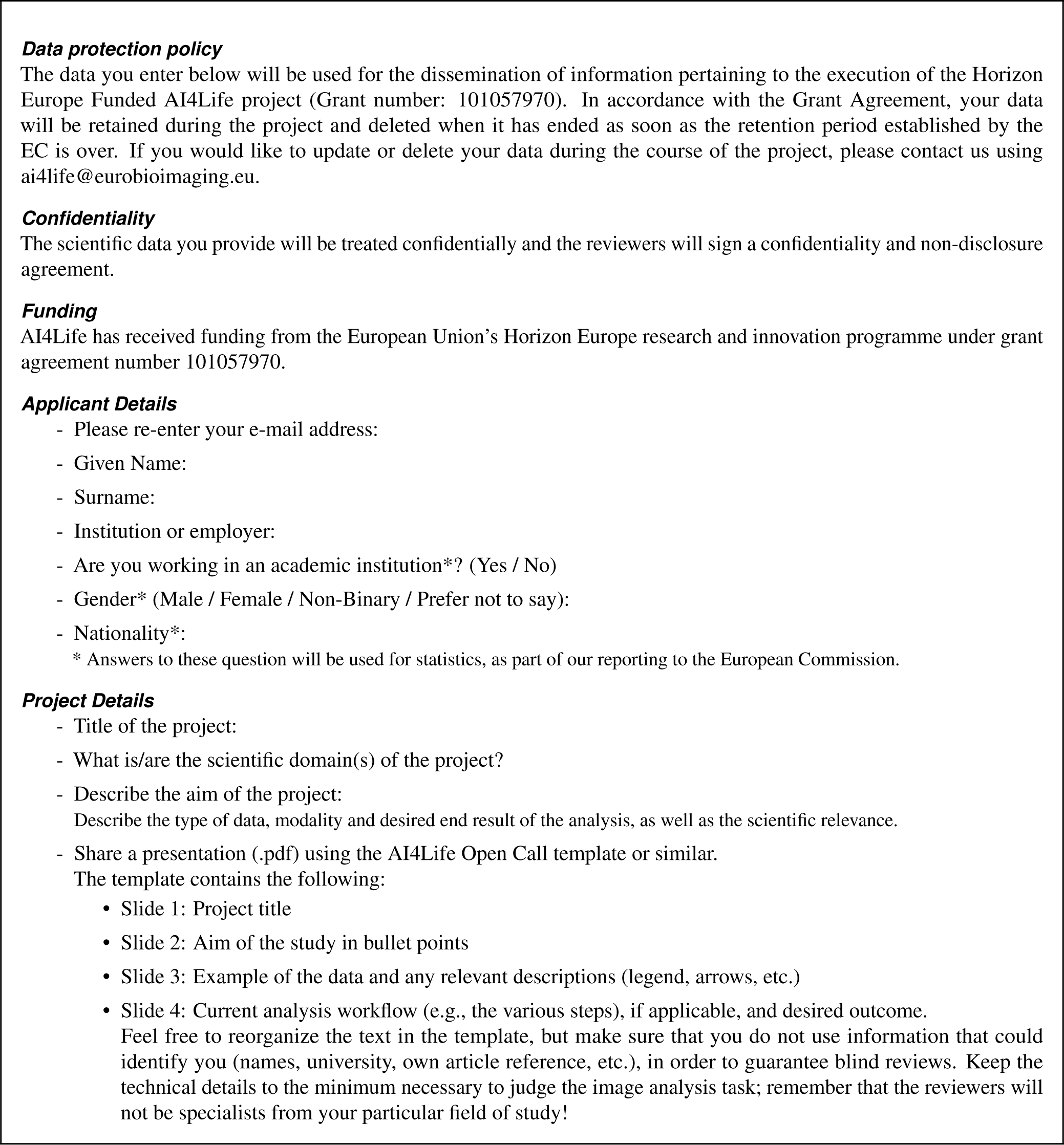
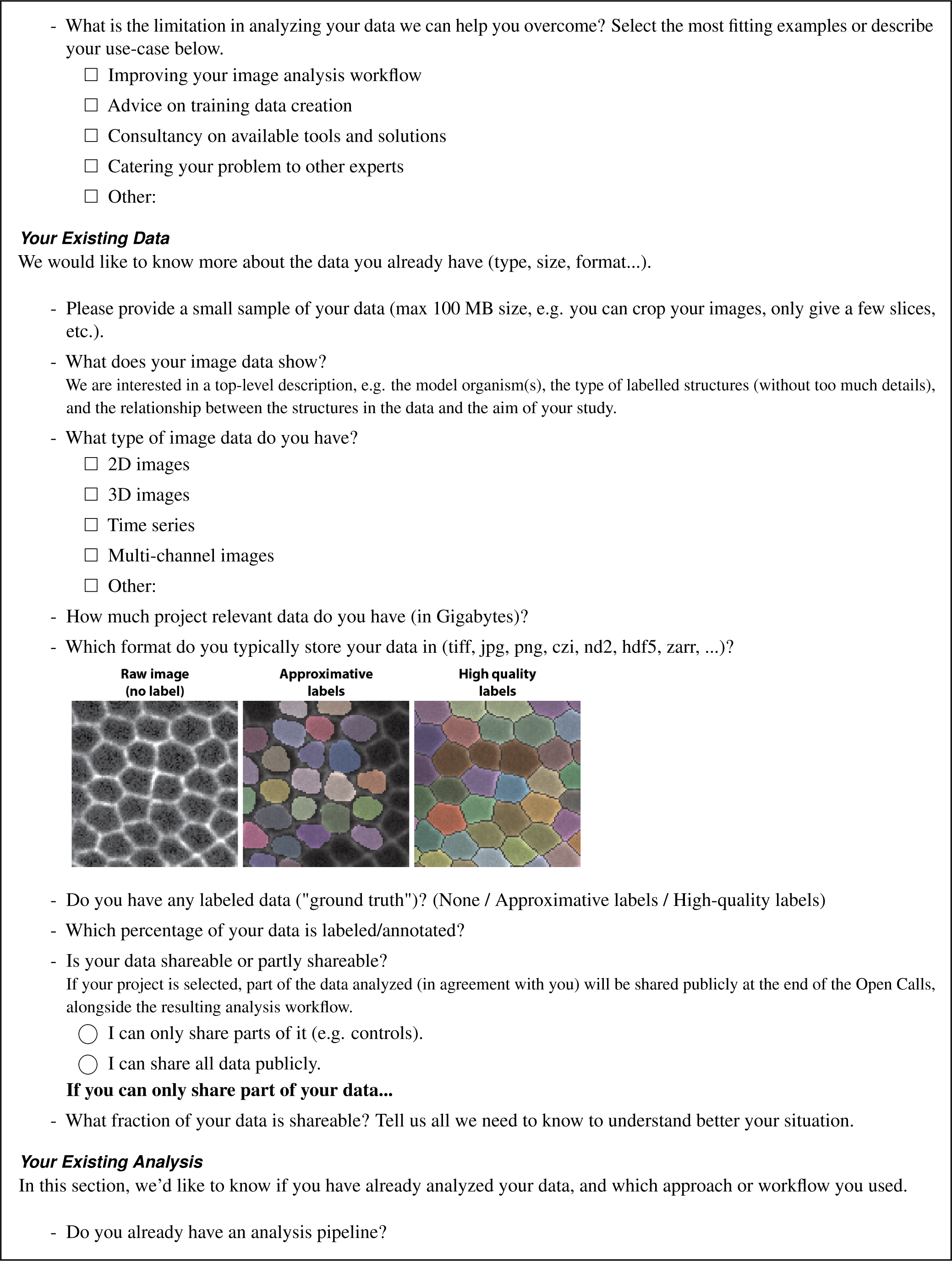
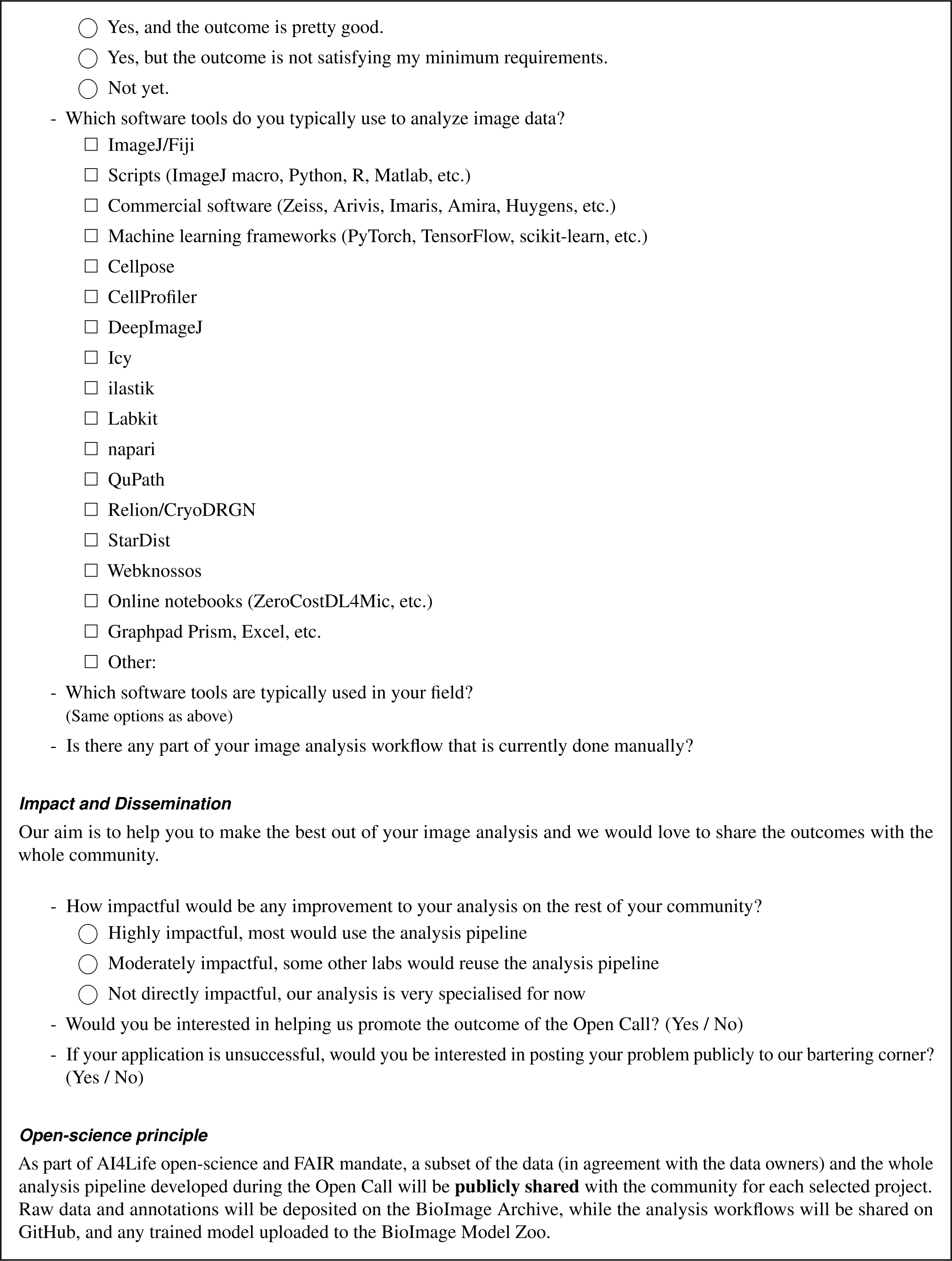
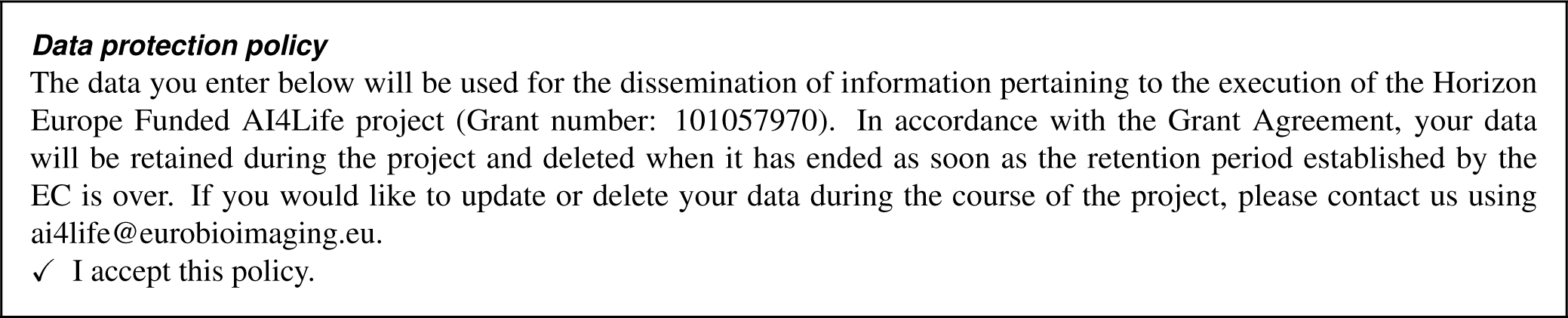

## A2 Appendix: OC 3 review form

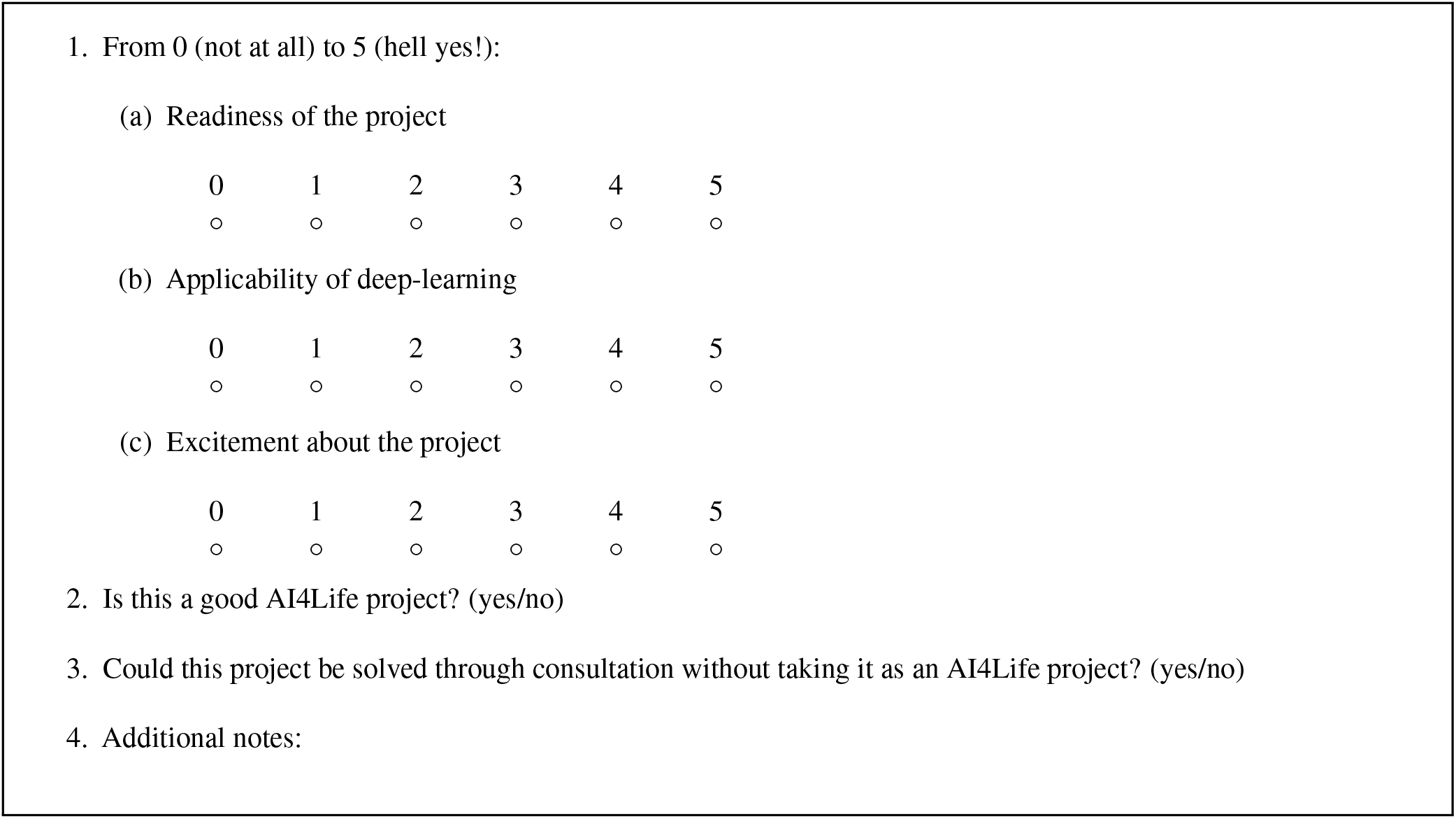

## A3 Appendix: Open Calls projects

**Table S8:** Selected projects in OC 1, 2 and 3.

| Open Calls | [Project ID] - Title | Description |
| --- | --- | --- |
| OC 1 | [3] - Treat CKD | The aim of the studying is to characterize the effect of Chronic Kidney Disease (CKD) on the density of collagen in specific tissues, such as the heart and kidney, using mouse and rat tissue slices. |
|  | [6] - Murine skeletal muscle analysis | The goal is to study the regulation of the metabolic properties in murine skeletal muscles, and the specific challenge is to segment, align and identify cells across individually acquired sections. |
|  | [10] - Image-guided gating strategy for image-enabled cell sorting of phytoplankton | The goal of this project iss to create an automated gating strategy for a cutting-edge commercial flow cytometer to sort phytoplankton from lab-grown cultures or field samples. |
|  | [11] - Segmentation and tracking of epithelial cells | The aim of this project is to develop a modular pipeline to segment and track epithelial cell nuclei in time-lapse fluorescence microscopy datasets, focusing on live imaging of dynamic epithelial monolayers. |
|  | [47] - Automated detection and quantification of senescent cells | Here, we aim at providing a semi-automated workflow to identify and quantify senescent cells in embryonic mouse neural tube tissue imaged by confocal microscopy. |
|  | [52] - Atlas of Symbiotic partnerships in plankton | The aim is to develop a segmentation workflow for identifying key organelles in FIB–SEM image volumes of microalgae engaged in symbiosis using limited annotations. |
|  | [66] - Automated and integrated cilia profiling | This project involves developing an automated detection and analysis of cilia length in cultured cells overexpressing motor proteins of the kif family. The goal is to correlate motor protein expression levels to cilia length. |
|  | [74] - Leaf tracker plant species proof | Developing a deep learning–based pipeline for instance segmentation and time-resolved tracking of plant leaves in RGB images across developmental stages. |

Table S8: Selected projects in OC 1, 2 and 3. (Continued)
| Open Calls | [Project ID] - Title | Description |
| --- | --- | --- |
| OC 2 | [11] - Automated species annotation in turbid waters | The project aim is to create an automated image-annotation pipeline to monitor the ecological status of vulnerable marine habitats by detecting various species in underwater photographic images. |
|  | [13] - Optimizing calcium image acquisition with machine learning denoising algorithms | The goal of the project is to benchmark various deep learning methods to denoise calcium imaging data acquired from in vivo brain recordings. |
|  | [14] - Diatom symbioses at planetary scale | The aim of the project is to identify symbiotic interactions between plankton species sampled from many ocean sites as part of the Tara Ocean project. |
|  | [21] - Automated segmentation of actin filaments in intact cells (EM) | Here, the aim is to segment actin filaments in cryo-ET tomograms, building a model that can generalize to future tomograms to produce quality segmentations. |
|  | [23] - Enhancing the effective resolution of intra-vital microscopy with digital expansion microscopy | Developing a deep learning pipeline capable of enhancing low-resolution in-vivo two-photon microscopy images to match the 4x resolution improvement obtained by expansion microscopy. |
|  | [26] - Ultrastructural protein mapping through Correlation of Light and Electron Microscopy | The project focuses on accurately aligning widefield fluorescence microscopy and transmission electron microscopy images to combine molecular and ultrastructural information for high-resolution analysis of endo-lysosomal alterations. |
|  | [38] - NeuroScan - A 3D human motor neuron disease platform for high throughput drug screening | The goal of the project is to segment nuclei and Islet1 positive nuclei in 3D neurospheres from their respective channels. |
| OC 3 | [3] - Improving nuclei segmentation using CellProfiler and StarDist | Enhancing DeepImageJ to run StarDist and other BioImage.io-compatible models directly within CellProfiler, ensuring seamless execution of advanced segmentation and feature-extraction routines. |
|  | [11] - 3D Matrix Motility Map (3DM<sup>3</sup>) | This project aims to analyze cancer cell dissemination in 3D extracellular matrices. |
|  | [14] - The speed of life in trees | The goal of this study is to automatically measure the fractional areas of vessels, axial and ray parenchyma, and fibers across the entire wood tissue slices and multiple species. |
|  | [17] - Imprints of Wind Disturbances on Wood Anatomy | To analyze differences in wood quality among trees experiencing varying degrees of wind disturbances. This requires an automatic segmentation of tree lumen and membrane in confocal microscopy. |
|  | [18] - Detection of Nuclear Pore Complexes with shape variability imaged by DNA PAINT | To identify nuclear pore complexes in DNA-PAINT images, including those that deviate from the canonical structure in terms of diameter, stoichiometry, and circularity. |
|  | [19] - Segmentation of sparse bacteria in human tissue | To segment and analyze bacteria clusters in human biopsies and characterize the biological microenvironment of niches (intra/extracellular, cell type, cluster size). |
|  | [26] - Automatic microtubule doublet picking in tomograms | To automatically detect and pick doublets with the correct polarity (i.e., all pointing the same) in Cryo-EM tomograms of cilia. |

**Table S9:**
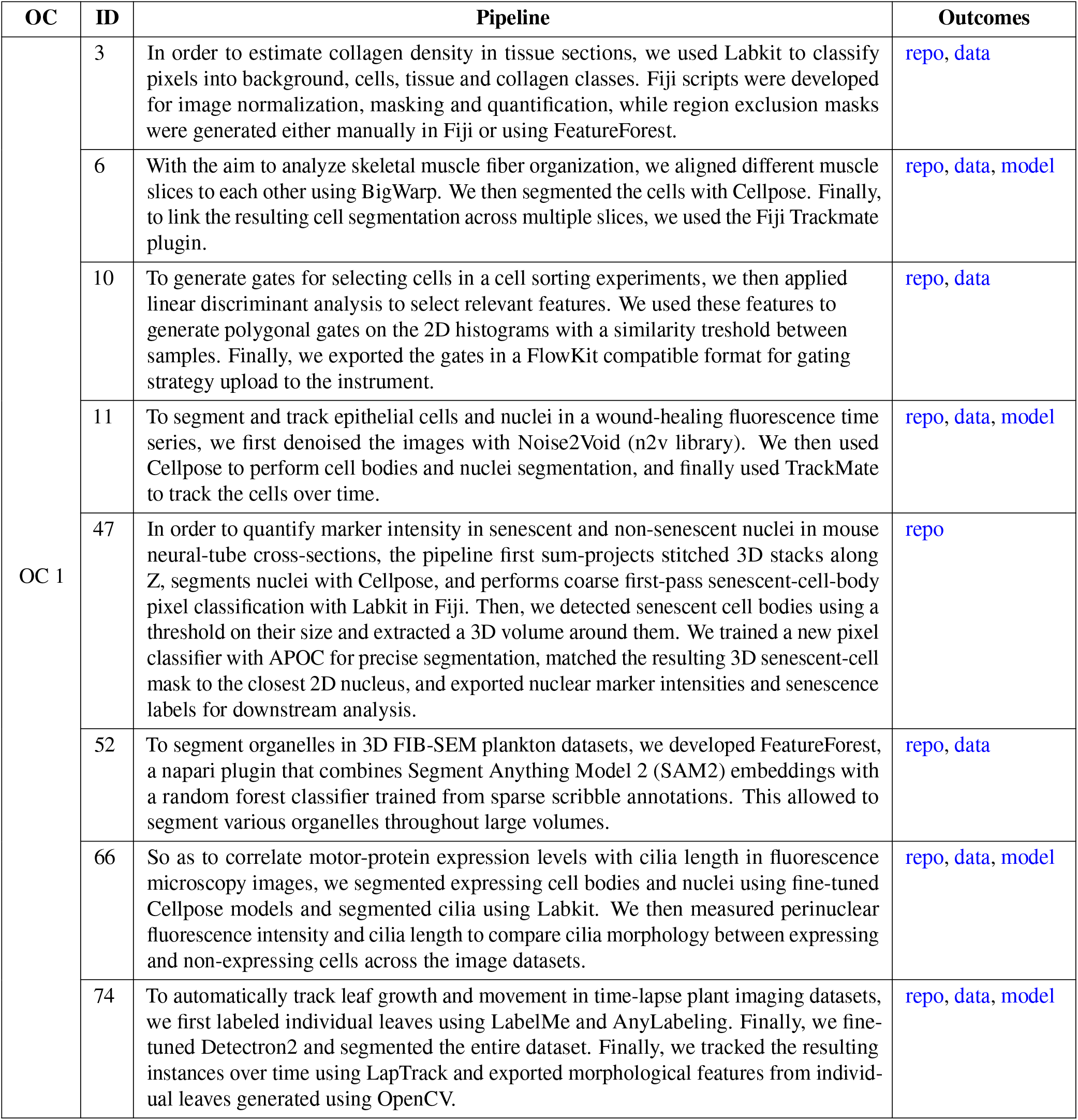
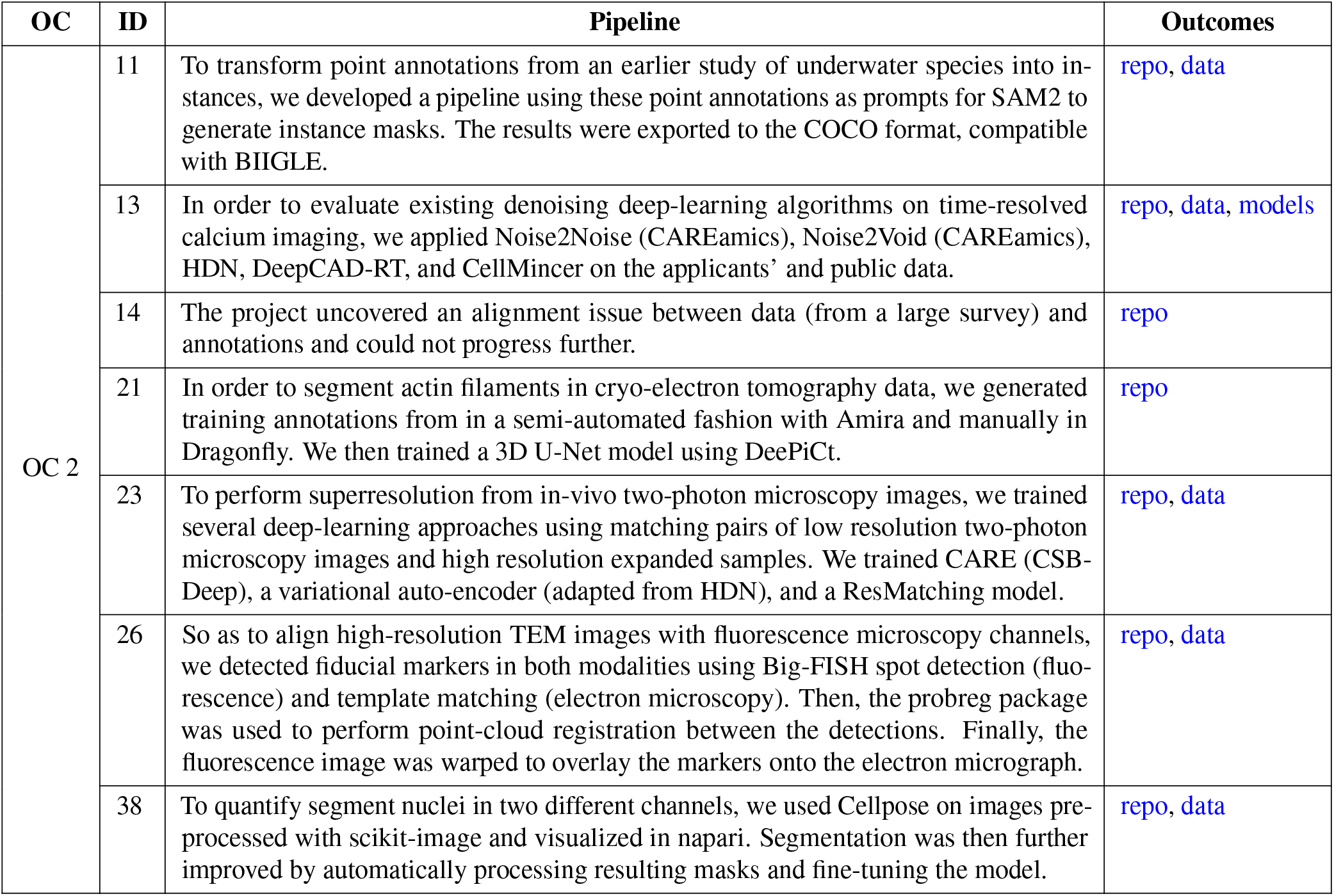
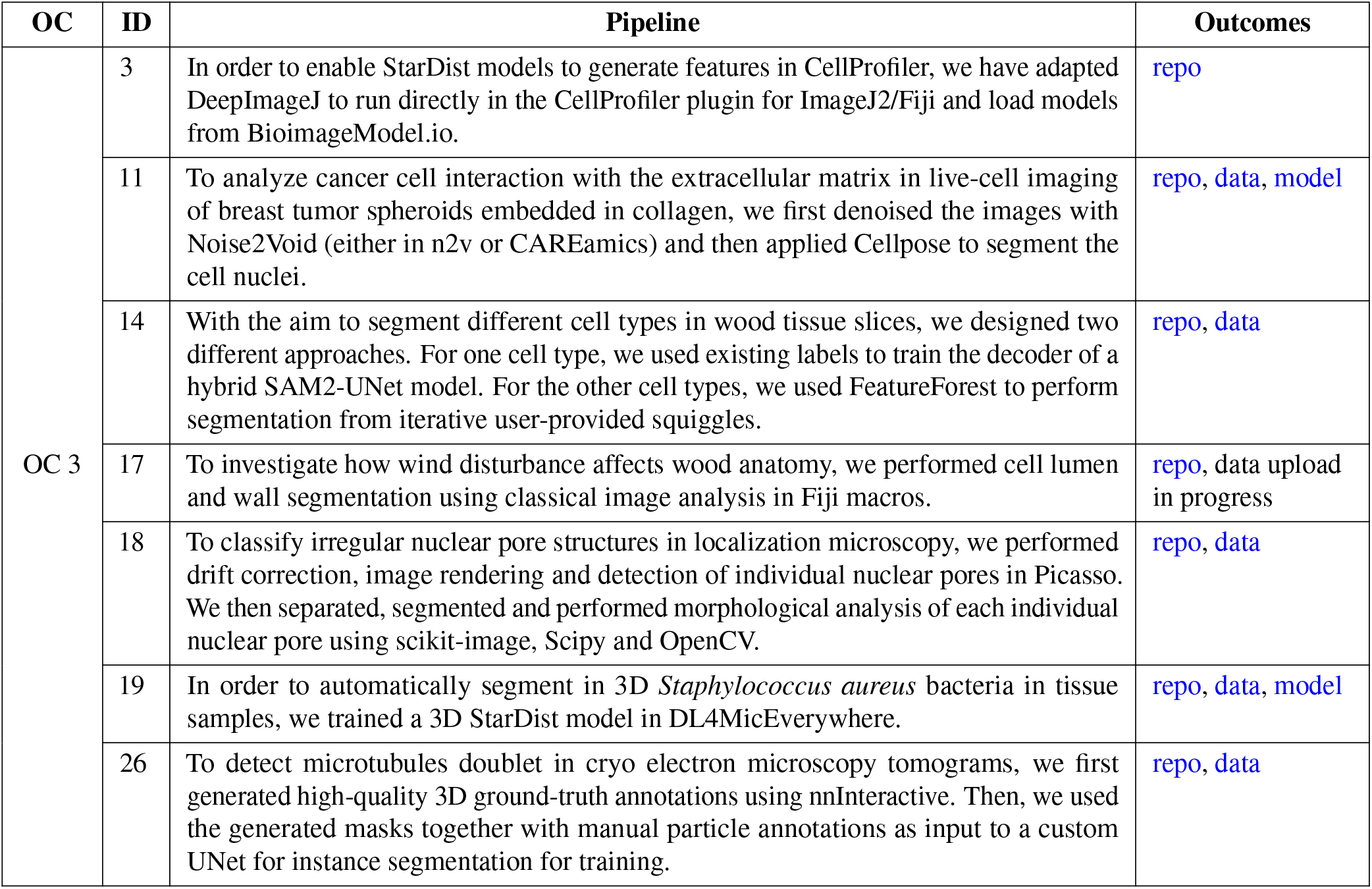
Project pipeline descriptions and outcomes.

